# Generation of iPSC lines with high cytogenetic stability from peripheral blood mononuclear cells (PBMCs)

**DOI:** 10.1101/2021.09.27.462082

**Authors:** Lindsay Panther, Loren Ornelas, Michelle R Jones, Andrew R. Gross, Emilda Gomez, Chunyan Liu, Benjamin Berman, Clive N. Svendsen, Dhruv Sareen

**Affiliations:** Board of Governors-Regenerative Medicine Institute, Cedars-Sinai Medical Center, Los Angeles, CA, 90048, USA; iPSC Core, The David and Janet Polak Foundation Stem Cell Core Laboratory; Center for Bioinformatics and Functional Genomics, Department of Biomedical Sciences, Cedars-Sinai Medical Center, Los Angeles, CA, 90048, USA.; Department of Biomedical Sciences, Cedars-Sinai Medical Center, Los Angeles, CA, 90048, USA.

**Author notes:** To whom correspondence should be addressed. Board of Governors-Regenerative Medicine Institute, Cedars-Sinai Medical Center, Los Angeles, CA, 90048. These authors contributed equally (Lindsay Panther, Loren Ornelas, Michelle R Jones). **Supplementary Tables can be viewed at:** https://docs.google.com/spreadsheets/d/1rboYasZNdsZ_tgMyhGQlqcxsH8CVBDhK/edit?usp=sharing&ouid=110828706727380725279&rtpof=true&sd=true.

**Keywords:** Human iPSCs, peripheral blood, skin fibroblasts, reprogramming, genetic integrity, genomic stability, karyotype, whole genome sequencing, regenerative medicine

## Abstract

The utility of human induced pluripotent stem cells (hiPSCs) is contingent upon genomic integrity and stability. Recurrent genomic aberrations have been observed in human iPSC lines upon long-term culture, ∼10-25% demonstrate karyotype abnormalities. We describe a new and reliable non-integrating episomal plasmid reprogramming method for fresh (unexpanded) peripheral blood mononuclear cells (PBMC) into iPSCs (PBMC-iPSCs). PBMC-iPSCs produced using this method have a superior chromosome-level karyotype stability rate (∼5% abnormality rate for all chromosomes; 2.8% for autosomes). After extended culture PBMC-iPSCs maintain a low rate of abnormalities (2% for autosomes). Deep coverage whole genome sequencing in a subset of PBMC-iPSC lines showed no shared single nucleotide polymorphisms (SNPs) or structural variants are introduced during reprogramming and maintenance of PBMC-iPSCs. iPSCs reprogrammed from unexpanded PBMCs have consistently high cytogenetic stability and minimal genomic aberrations, suggesting this method is highly suited for iPSCs in research and therapeutic clinical applications.

## INTRODUCTION

Human embryonic stem cells (hESCs), discovered in 1998 (Thomson *et al*., 1998), had promise to be a universal source of cell replacement therapies for many degenerative diseases. However, their embryonic origin raises ethical concerns, and they are limited in the generation of a wide array of disease and patient-specific cell lines. Over the past decade, human induced pluripotent stem cells (hiPSCs) have emerged as promising biological tools for the study of human development, human disease modeling, cell fate decisions, tissue regeneration, and novel therapeutic drugs(Ebert and Svendsen, 2010; Zeltner and Studer, 2015; Soria-Valles and López-Otín, 2016). These cells provide a continuous supply of well-characterized stem and progenitor cells from patients with disparate genotypes. Critically, hiPSCs provide an autologous source of cells and hence avoid immune rejection, giving them enormous potential as a cell therapy for regenerative medicine(Kamao *et al*., 2014; Masayo Takahashi, 2016).

Maintaining genomic integrity and stability of hiPSC lines is imperative for reliable disease modeling and safe clinical applications of stem cells in regenerative therapies. Aberrant cytogenetic errors that arise during reprogramming of somatic cells as well as during expansion and maintenance of hiPSCs impact the accuracy of *in vitro* disease modelling and, more crucially, the *in vivo* utility of iPSCs for regenerative medicine. It is important that iPSCs in clinical use are free from cancer-associated genomic aberrations, especially given that several studies have reported chromosomal aneuploidy, translocations, duplications and deletions, and point mutations in iPSCs (Mayshar *et al*., 2010; Gore, Li, Fung, Jessica E. Young, *et al*., 2011; Taapken *et al*., 2011; Hussein *et al*., 2011; Laurent *et al*., 2011; Martins-Taylor *et al*., 2011; Young *et al*., 2012; Ma *et al*., 2014; Garber, 2015; Salomonis *et al*., 2016; Kilpinen *et al*., 2017; Lo Sardo *et al*., 2017; Merkle *et al*., 2017).

Various reprogramming approaches have been utilized, including genome-integrating DNA elements such as lentiviral, retroviral and transposon vectors, and the newer generation of non-integrating methods such as adenoviral and Sendai viral vectors, protein transduction, small molecules, synthetic mRNA, miRNA and episomal plasmids (Schlaeger *et al*., 2015). However, many of the non-integrating reprogramming methods suffer from low reprogramming efficiency and some require serial transgene or protein deliveries into the cell. Regardless of the chosen reprogramming method, it is advantageous for clinical applications that the reprogrammed iPSCs maintain original genomic integrity and do not carry genomic integrations of reprogramming factors and integrated vector sequences.

The most common source for human iPSC derivation has been dermal skin fibroblasts, either from fresh skin biopsies or stored fibroblast lines in cell repositories. However, the requirement for skin biopsies to expand and bank fibroblast cells for several passages represents an impediment that must be overcome to make iPSC technology broadly applicable. The low-level background mutations in the parental fibroblasts are also a likely confounding factor in the long-term stability of isolated iPSC clones post reprogramming (Young *et al*., 2012). Furthermore, a skin biopsy typically requires a 3-4 mm skin punch and local anesthetics. Instead, peripheral blood can circumvent issues with the common fibroblast cell source and is therefore a promising source of patient tissue for reprogramming. Peripheral blood is not only minimally invasive to collect, it is often accessible through the large numbers of frozen patient samples stored in blood biorepositories. After a patient blood draw, reprogramming PBMCs into iPSCs later requires an intermediate PBMC cryopreservation step to reliably recover viable blood cells. This process provides great flexibility because select cohorts of cryopreserved PBMCs from patients can be converted to iPSCs when further patient genotype-phenotype information is available. A cryopreservation step for PBMCs thus allows for iPSC “future-proofing”.

Increased sub-chromosomal copy number variations (CNVs) have been reported in iPSCs (Gore, Li, Fung, Jessica E. Young, *et al*., 2011; Hussein *et al*., 2011; Laurent *et al*., 2011; Martins-Taylor *et al*., 2011; Taapken *et al*., 2011), including deletions associated with tumor-suppressor genes, and duplications of oncogenes, with differences in early, intermediate or late passage numbers. In these studies, most genomic abnormalities occurred in iPSC lines generated from dermal fibroblasts using integrating reprogramming methods. To date, however, no systematic reports of recurrent sub-chromosomal abnormalities specific to large numbers of fibroblast-, epithelial- or blood-derived hiPSC lines have been described, in which similar episomal non-integrating reprogramming and stem cell culture methods are used.

In this study, we describe a reliable method to efficiently reprogram peripheral mononuclear blood cells (PBMCs) into iPSCs (PBMC-iPSCs) from both a lymphoid T cell and a myeloid non-T cell population. These PBMC-iPSC lines show far greater genetic stability when compared to iPSCs derived from previously expanded cell types, including dermal fibroblasts (fib-iPSCs), lymphoblastoid cell lines (LCLs), epithelial or adipose stem cells that were obtained from public donor cell repositories or research laboratories. Thus, the reprogramming method reported here for unexpanded and cryopreserved PBMCs results in more genetically stable iPSCs.

## RESULTS

### Cryopreserved PBMCs Isolated from Blood are Reliably Reprogrammed to iPSCs

Human iPSCs can be generated from freshly isolated unexpanded PBMCs using episomal plasmids (Okita *et al*., 2013). However, there are no descriptions for reprogramming cryopreserved PBMCs to iPSCs. When we reprogrammed cryopreserved PBMCs to iPSCs from multiple individuals with episomal plasmids expressing *POU5F1*, *SOX2*, *KLF4*, *LIN28*, *L-MYC*, and *TP53* shRNA (previously described as the 4p method (Okita *et al*., 2013)), we were either unsuccessful or observed significant variability in isolating identifiable iPSC clones even after 35–40 days (**Supplementary Table 1**). To generate sufficient cell numbers for reprogramming isolated CD34^+^ progenitor cells need to be enriched and expanded in culture with complex and expensive protocols (Loh *et al*., 2009; Ban *et al*., 2011; Okita *et al*., 2013; Schlaeger *et al*., 2015), highlighting the need to simplify this process and avoid expansion of somatic cells prior to reprogramming. We instead isolated and cryopreserved unexpanded PBMCs for reprogramming to iPSCs as a source of somatic cells to provide the fastest and most cost-effective procedure from large, multi-subject cohorts. Using the 4p method the T reprogramming protocol was successful with only 14.3% of PBMC samples, while the non-T protocol used with cryopreserved PBMCs did not result in any clonal iPSC lines (**Supplementary Table 1**).

To increase the efficiency of PBMC reprogramming we utilized, (a) an additional episomal plasmid containing SV40 large T antigen (SV40LT) in specific stoichiometry to minimize PBMC cell death, which we have termed the 5p method, and (b) a defined E7 reprogramming media to promote high surface attachment of the nucleofected PBMCs (Fig. 1a). This novel protocol resulted in successful and efficient generation of multiple adherent PBMC-iPSC clones that could be mechanically isolated and scaled up for expansion within 25–35 days post-nucleofection (Fig. 1a). Importantly, the success rate and the efficiencies in reprogramming multiple subject cryopreserved PBMCs from unaffected controls or diseased patients was significantly greater at 86% for T cell and 74% for non-T cell types when using the 5p method, in contrast to between 0 and 14% for the 4p method (Fig. 1b). This effect was observed when culturing with either mouse embryonic fibroblast feeders (MEFs) or xenobiotic-free components using recombinant human laminin 521 substrates and chemically defined reprogramming media (E7) (Chen *et al*., 2011) (**Supplementary Table 2**). Collectively, these results show that the 5p method, when compared to the 4p method, significantly enhanced reprogramming success of PBMCs (both T and non-T cells) (Figs. 1b and c) and provides a valuable protocol for reliable reprogramming of cryopreserved PBMCs.

**Figure 1.**
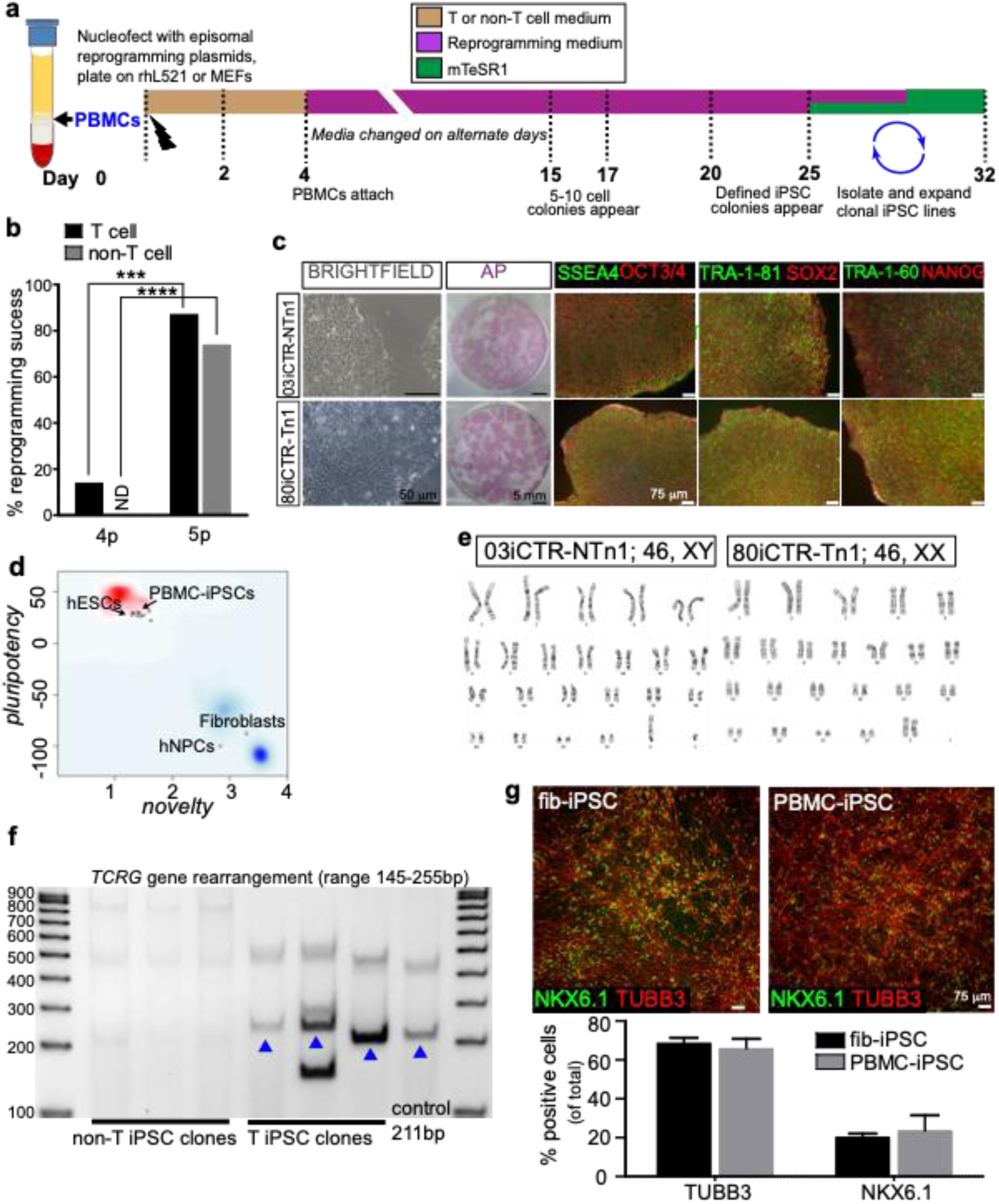
Reliable Episomal Reprogramming of PBMCs into iPSCs. (a) Schematic of the episomal reprogramming process and timeline of iPSC generation from PBMCs using the 5p method. (b) Percent of iPSC reprogramming success for cryopreserved PBMCs by the 5p method when compared to the published 4p method for T cell and non-T cell lines. (c) Representative images of brightfield, alkaline phosphatase and ICC stains of key pluripotent markers for a T-cell (80iCTR) and non-T-cell line (03iCTR). (d) Pluripotency and novelty scores of reprogrammed lines according to the microarray-based PluriTest analysis. (e) Healthy representative G-band karyotypes of a T cell (80iCTR) and non-T cell (03iCTR) line.(g) T cell receptor gene rearrangement assay shows different sets of PCR primers were used to detect *TCRG* gene rearrangements occurring in the PBMC-iPSCs derived from the T-cell method, while PCR products in the 145-255 bp range were not detectable in the non-T cell iPSC clones. (h) Directed neuronal differentiation of representative fibroblast-derived (*n* = 26) and PBMC-derived iPSCs (*n* = 20) showing equivalent numbers of TUBB3+ (β_3_-tubulin+) and NKX6.1+ neuronal cells.

Each PBMC-iPSC line exhibited typical PSC characteristics, including tightly packed colonies, high cell nuclear-cytoplasmic ratio, robust alkaline phosphatase activity, and expression of multiple pluripotency antigens. These features are shown in representative reprogrammed iPSC colonies from PBMCs of two healthy volunteers (non-T cell source 03iCTR-NTn1 and T-cell source 80iCTR-Tn1) (Fig. 1c). The PBMC-iPSCs passed pluripotency quality control metrics determined by the PluriTest assay(Müller *et al*., 2011), demonstrating that the PBMC-iPSC transcription profile was analogous to well established hESCs and fib-iPSCs, but not differentiated fibroblasts and neural progenitor cells (Fig. 1d). PBMC-iPSCs maintained a normal G-band karyotype (Fig. 1e) and were confirmed to be clonal derivatives of either T cells or non-T cells from the PBMCs based on the T cell receptor (TCR)-β and –γ, gene rearrangement/clonality assays (Fig. 1f). The tri-lineage potential of PBMC-iPSCs was demonstrated by spontaneous embryoid body formation and by measuring germ-layer specific gene expression profile by the TaqMan Scorecard assay (Supplementary Fig. 1). The genomic transgene-free status of PBMC-iPSCs was confirmed by, (1) demonstration that the EBNA plasmid-related latency element was eventually eliminated from the established PBMC-iPSCs and not detected in the genomic DNA (Supplementary Fig. 2a); and (2) the expression of endogenous pluripotency genes and absence of any exogenous reprogramming transgenes using RT-qPCR (Supplementary Fig. 2b). Further, the SV40LT factor that is a component of the 5p reprogramming cocktail was undetectable at the genomic DNA level (Supplementary Fig. 3a) and not expressed by the reprogrammed PBMC-iPSC lines (Supplementary Fig. 3b).

### PBMC-iPSCs have Equivalent Neuronal Differentiation as Expanded Fib-iPSCs

While many early reports suggested that newly reprogrammed iPSCs retain some epigenetic memory of their parental donor cell type (Barrero, Boué and Izpisúa Belmonte, 2010; Jaenisch *et al*., 2010; Lister *et al*., 2011; Ohi *et al*., 2011), recent reports suggest that hiPSCs, regardless of the source tissue, ultimately lose most of their gene expression and epigenetic profiles related to the original cell source following expansion (Sareen *et al*., 2014; Kyttälä *et al*., 2016). However, it still remains unclear whether blood-derived iPSCs can differentiate as efficiently as fibroblast-derived iPSCs into various cell types, possibly due to a stronger retention of epigenetic memory in blood-sourced iPSCs compared to other donor cells (Jaenisch *et al*., 2010; Kim *et al*., 2011). We addressed this by directing blood or fibroblast-derived iPSCs, both of mesodermal origin, towards neuroectoderm, a different germ layer. We performed neural ectoderm differentiation from iPSC lines derived from fibroblast (*n* = 26) and PBMC lines (*n* = 20) of both healthy subjects and diseased patients. Immunocytochemistry and quantification for neural ectoderm markers of TUBB3 (β_3_-tubulin) and NKX6.1 demonstrated that neuronal differentiation occurred at a similar efficiency between expanded fib-iPSC and unexpanded PBMC-iPSCs (Fig. 1g), further demonstrating the utility of unexpanded PBMC-iPSCs.

### PBMC-iPSCs Reprogrammed with 5p Maintain a Significantly More Stable Karyotype

Recurrent chromosomal abnormalities have been described previously for hESC lines (Baker *et al*., 2007; Spits *et al*., 2008; Taapken *et al*., 2011). Some reports have also chronicled common chromosomal aberrations for hiPSC lines; however, these did not methodically account for variability in the source tissues, their levels of expansion prior to reprogramming, methods of reprogramming and cell culture. To our knowledge, there have been no systematic studies describing cytogenetic analyses comparing aberration frequency between iPSCs derived from expanded cell sources and unexpanded cell types such as PBMCs isolated from whole blood. We performed in-depth cytogenetic analysis on iPSC lines derived from 1,028 unique donors of fibroblasts (n=98), LCLs (n=54), epithelial (n=15), adipose (n=1) and PBMCs (n=860) sourced from diverse laboratories and public repositories. Multiple clones were derived from each of these parent lines, yielding 402 i PSC lines generated from expanded cell lines and 1,063 iPSC lines generated from PBMCs. We performed repeated karyotype analysis on a subset of iPSC lines, yielding a total of 526 karyotypes of iPSCs from expanded cells and 1,877 karyotypes from unexpanded PBMCs. In total, G-band karyotype analysis was completed on 2,443 samples derived from cultured sources including fibroblasts (n=398), LCLs (n=122), epithelial (n=42), adipose (n=4), and PBMC (n=1877). All fibroblast, LCLs, epithelial, adipose and PBMC derived iPSCs assessed were reprogrammed using similar non-integrating episomal reprogramming methods. The major difference between fibroblast and PBMC-reprogramming reported here was the number of reprogramming factor plasmids used; 3p (Okita *et al*., 2011) was used to derive lines from fibroblasts and 4p or 5p for PBMC derived lines. After successful reprograming all iPSC lines analyzed and reported here were maintained using feeder-free Matrigel®/mTeSR1™ cell culture methods and passaged using mechanical StemPro® EZPassage™ tool.

Our data reveal remarkably lower incidence of chromosomal aberrations in unexpanded PBMC-iPSCs. Abnormal karyotypes were observed in 96 of 526 (18.3%) cultures from clonally independent expanded source tissue - derived iPSC lines, derived from 168 unique donors (Fig. 2). Typically, the expanded cell types were expanded for 4-12 passages prior to reprogramming. In stark contrast, only 95 in 1,877 (5.1%) cultures of clonal human PBMC-iPSC lines derived from 860 unique donors displayed any karyotype abnormalities, and the majority of observed abnormalities were low-frequency. Recurrent aberrations were represented in a significantly greater degree in expanded source tissue-derived-iPSCs compared to the unexpanded PBMC-iPSCs (Fig. 2). This remarkable cytogenetic stability is unique to unexpanded PBMCs reprogrammed with the new 5p method. The most common abnormality in all cases was X-chromosome monosomy (45,X mosaicism), which is common and likely a harmless finding in blood cultures from normal women, possibly due to aging and errors occurring in cell divisions during which the inactivated X-chromosome is sometimes lost (Surrallés *et al*., 1999). 45,X karyotype cells are also seen in the blood of normal males though at a lower rate than females, where about 2% of lymphocytes in 30 year old males are missing a Y chromosome. An average of 10% of cells showed sex chromosome loss in 5 centenarian males (Bukvic *et al*., 2001). Excluding all sex chromosome abnormalities the karyotype abnormality rate for unexpanded PBMC-iPSCs reduces to 2.8% compared to 15.6% for expanded source-derived iPSCs. A complete list of all karyotypes, the respective donor and results of karyotype analysis can be found in **Supplementary Table 3**.

**Figure 2.**
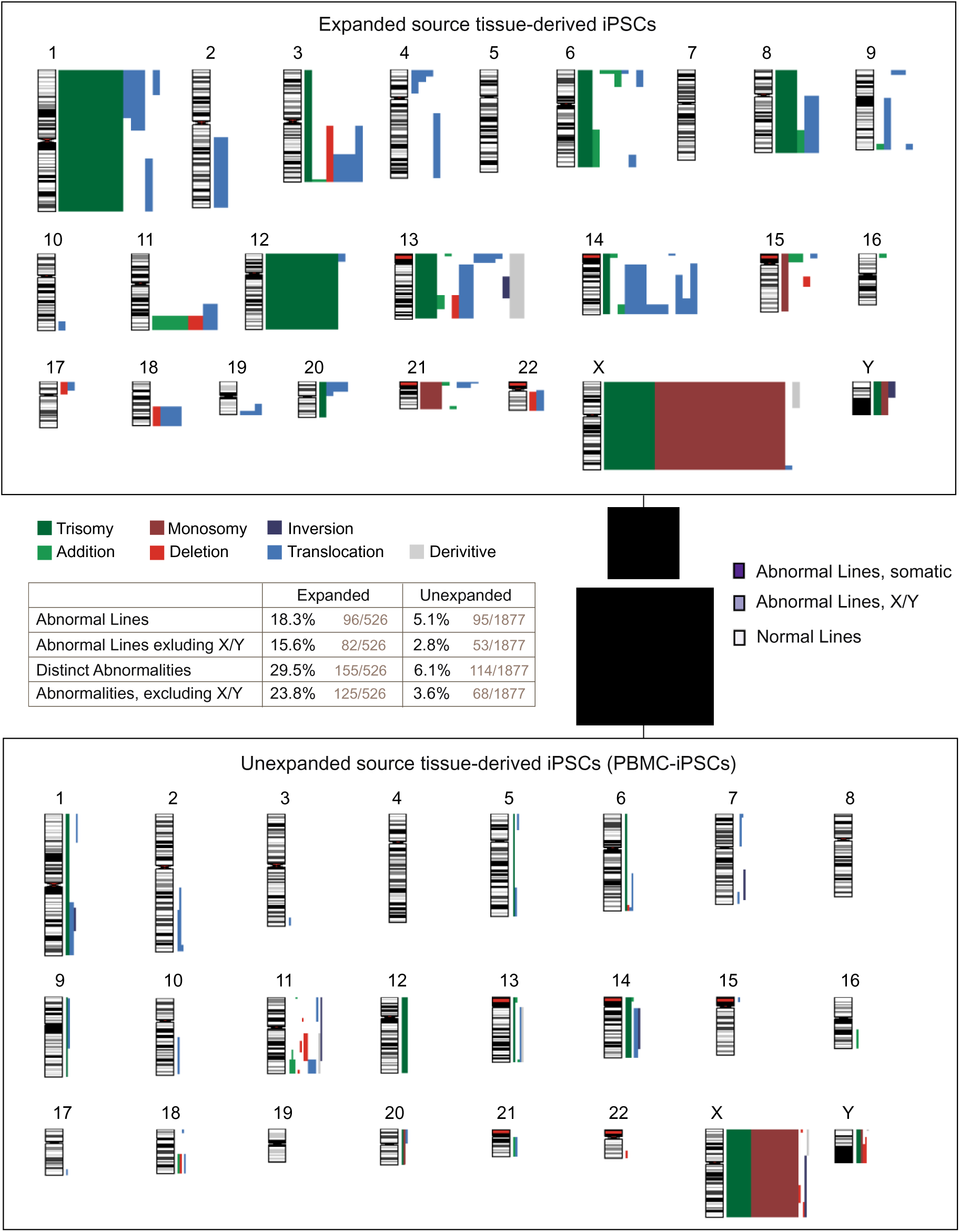
PBMC-iPSCs Derived by 5p Method Exhibit Stable Karyotypes. All detected karyotypic abnormalities were plotted on their respective loci for expanded (top) and unexpanded cells (bottom). Because 3.5x as many unexpanded lines were analyzed as expanded lines 1877 vs. 526), the bar widths were scaled accordingly. The fraction of lines with a normal karyotype for each group was plotted in two pie charts, which were also scaled to visually communicate the sample number for reach group. Among the 526 expanded cultures, 18.3% (96) were abnormal, and 15.6% (82) were abnormal discounting abnormalities on sex chromosomes. Among the 1877 unexpanded cultures, 5.1% (95) were abnormal, and 2.8% (53) when discounting sex chromosomes. Expanded cultures exhibited a total of 155 distinct abnormalities, of which 125 were on somatic chromosomes. Unexpanded cultures exhibited 114 abnormalities, of which 68 were on somatic chromosomes.

The most frequent karyotype abnormalities in expanded-iPSCs were observed in chromosomes X (17.4%), 13 (9.7%), 1 (9.0%), 14 (8.4%), and 12 (7.1%) (Table 1). To a lesser extent karyotype changes were also observed in chromosomes 11, 3, and 21. Interestingly, karyotype aberrations were never detected on chromosomes 5 and 7 in expanded iPSCs. Of the aneuploidies, chromosomal gains (trisomy or duplications) were most commonly observed in 12 (17.5%), 1 (15.8%), and X (10.5%), and chromosomal losses were repeatedly observed in chromosomes X (60%) and 21 (10%). Structural rearrangements, including translocations, inversions and derivative chromosomes, were recurrent in chromosomes 14 (15.6%), 13 (14.1%) and 1 (7.8%) in expanded-iPSCs. In stark contrast, in unexpanded PBMC-iPSC lines gain or loss of sex chromosomes X and Y were the most common aneuploidy, while rearrangements in chromosome 11 were observed in the few lines with karyotype abnormalities (Table 2).

**Table 1.**
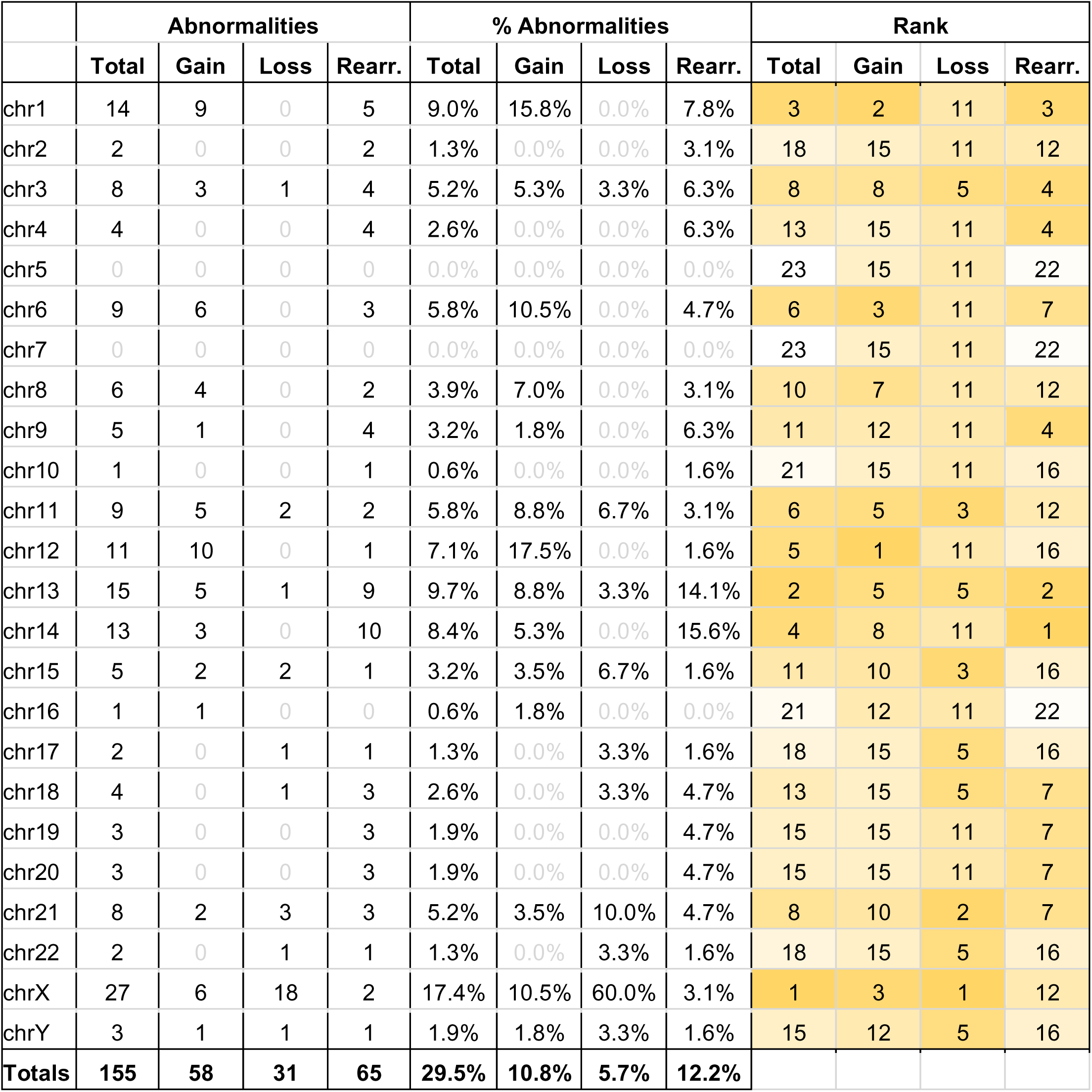
Frequency of karyotype abnormalities in expanded source tissue-derived iPSCs observed by G-band karyotyping. Cells shaded in orange depict the highest ranked percent karyotype abnormality for each of the categories of total chromosomal abnormalities, which include gain, loss or structural rearrangements per chromosome. Rearr. – Structural Rearrangements including translocations, inversions and derivatives.

**Table 2.** Frequency of the different karyotype abnormalities in unexpanded PBMC-derived iPSCs observed by G-band karyotyping. Cells shaded in orange depict the highest ranked percent karyotype abnormality for each of the categories of total chromosomal abnormalities, which include gain, loss or structural rearrangements per chromosome. Rearr. – Structural Rearrangements including translocations, inversions and derivatives.

|  | Abnormalities |  |  |  | % Abnormalities |  |  |  | Rank |  |  |  |
| --- | --- | --- | --- | --- | --- | --- | --- | --- | --- | --- | --- | --- |
|  | Total | Gain | Loss | Rearr. | Total | Gain | Loss | Rearr. | Total | Gain | Loss | Rearr. |
| chr1 | 14 | 9 | 0 | 5 | 9.0% | 15.8% | 0.0% | 7.8% | 3 | 2 | 11 | 3 |
| chr2 | 2 | 0 | 0 | 2 | 1.3% | 0.0% | 0.0% | 3.1% | 18 | 15 | 11 | 12 |
| chr3 | 8 | 3 | 1 | 4 | 5.2% | 5.3% | 3.3% | 6.3% | 8 | 8 | 5 | 4 |
| chr4 | 4 | 0 | 0 | 4 | 2.6% | 0.0% | 0.0% | 6.3% | 13 | 15 | 11 | 4 |
| chr5 | 0 | 0 | 0 | 0 | 0.0% | 0.0% | 0.0% | 0.0% | 23 | 15 | 11 | 22 |
| chr6 | 9 | 6 | 0 | 3 | 5.8% | 10.5% | 0.0% | 4.7% | 6 | 3 | 11 | 7 |
| chr7 | 0 | 0 | 0 | 0 | 0.0% | 0.0% | 0.0% | 0.0% | 23 | 15 | 11 | 22 |
| chr8 | 6 | 4 | 0 | 2 | 3.9% | 7.0% | 0.0% | 3.1% | 10 | 7 | 11 | 12 |
| chr9 | 5 | 1 | 0 | 4 | 3.2% | 1.8% | 0.0% | 6.3% | 11 | 12 | 11 | 4 |
| chr10 | 1 | 0 | 0 | 1 | 0.6% | 0.0% | 0.0% | 1.6% | 21 | 15 | 11 | 16 |
| chr11 | 9 | 5 | 2 | 2 | 5.8% | 8.8% | 6.7% | 3.1% | 6 | 5 | 3 | 12 |
| chr12 | 11 | 10 | 0 | 1 | 7.1% | 17.5% | 0.0% | 1.6% | 5 | 1 | 11 | 16 |
| chr13 | 15 | 5 | 1 | 9 | 9.7% | 8.8% | 3.3% | 14.1% | 2 | 5 | 5 | 2 |
| chr14 | 13 | 3 | 0 | 10 | 8.4% | 5.3% | 0.0% | 15.6% | 4 | 8 | 11 | 1 |
| chr15 | 5 | 2 | 2 | 1 | 3.2% | 3.5% | 6.7% | 1.6% | 11 | 10 | 3 | 16 |
| chr16 | 1 | 1 | 0 | 0 | 0.6% | 1.8% | 0.0% | 0.0% | 21 | 12 | 11 | 22 |
| chr17 | 2 | 0 | 1 | 1 | 1.3% | 0.0% | 3.3% | 1.6% | 18 | 15 | 5 | 16 |
| chr18 | 4 | 0 | 1 | 3 | 2.6% | 0.0% | 3.3% | 4.7% | 13 | 15 | 5 | 7 |
| chr19 | 3 | 0 | 0 | 3 | 1.9% | 0.0% | 0.0% | 4.7% | 15 | 15 | 11 | 7 |
| chr20 | 3 | 0 | 0 | 3 | 1.9% | 0.0% | 0.0% | 4.7% | 15 | 15 | 11 | 7 |
| chr21 | 8 | 2 | 3 | 3 | 5.2% | 3.5% | 10.0% | 4.7% | 8 | 10 | 2 | 7 |
| chr22 | 2 | 0 | 1 | 1 | 1.3% | 0.0% | 3.3% | 1.6% | 18 | 15 | 5 | 16 |
| chrX | 27 | 6 | 18 | 2 | 17.4% | 10.5% | 60.0% | 3.1% | 1 | 3 | 1 | 12 |
| chrY | 3 | 1 | 1 | 1 | 1.9% | 1.8% | 3.3% | 1.6% | 15 | 12 | 5 | 16 |
| **Totals** | **155** | **58** | **31** | **65** | **29.5%** | **10.8%** | **5.7%** | **12.2%** |  |  |  |  |

### Enhanced Karyotype Stability of PBMC-iPSCs is Independent of Donor Age and Passage Number

Unlike whole blood, other somatic cell types including skin fibroblasts, epithelial and adipose cells derived from tissue biopsies require a certain level of expansion in culture prior to reprogramming, which is akin to further aging in culture. Some of these tissues such as skin may also be routinely exposed to external environmental elements like potential sun damage. Given this, we posited that the donor age might be a contributing factor that impacts the cytogenetic instability of expanded tissue derived-iPSCs. G-band karyotype analysis was used to assess this possibility. There was a significant increase in the frequency of cytogenetic abnormalities of expanded iPSCs with increasing age, especially above the donor age of 80. Among expanded lines 16.5% of the 363 lines from donors ≤80 years of age exhibited abnormalities, compared to 40% of the 30 lines from donors >80 years of age (P-value=0.0014, Z-Score=3.193). Conversely, among unexpanded lines 5.3% of the 1,595 lines from donors ≤80 years of age exhibited abnormalities versus 0% of the 23 lines from donors >80 years of age (P-value=0.26, Z-score=1.13). There was no significant trend observed between the donor ages of 0-60. In contrast, these results confirmed that unexpanded PBMC-iPSCs had significantly greater karyotype stability over expanded-iPSCs in every donor age group. Donor age was not an important factor in impacting karyotype abnormality frequencies for unexpanded PBMC-derived iPSCs (Fig. 3a) (**Supplementary Table 4**).

**Figure 3.**
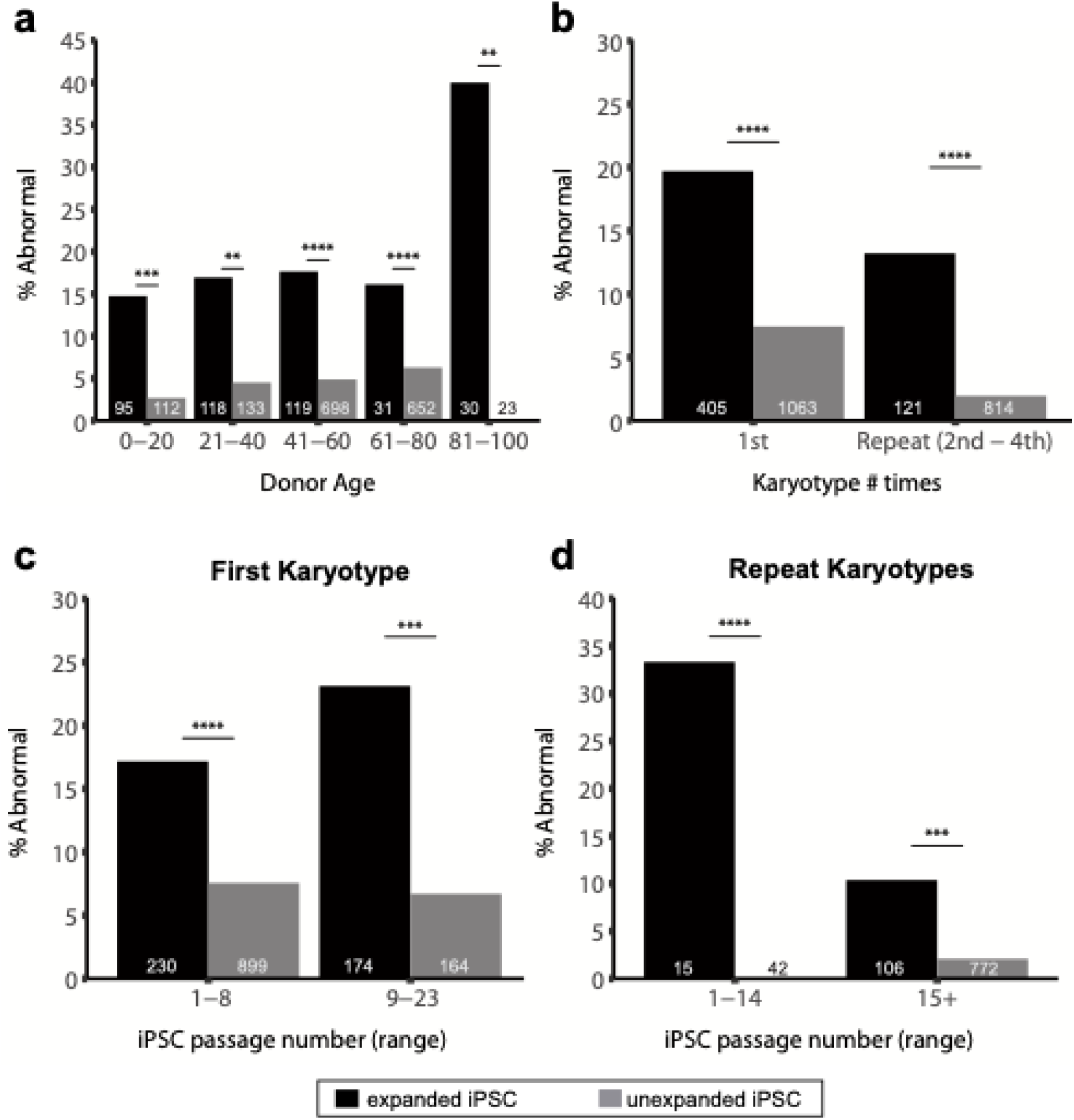
The PBMC-iPSCs Maintain Remarkably More Stable Karyotypes Over Extended Culture, Independent of Donor Age. For all subfigures the total number of samples, n, is displayed at the base of each bar. (a) The fraction of abnormal karyotypes within five age groups. (b) A comparison of the percent of karyotypes which were abnormal on the first karyotype and on subsequent follow-up karyotypes for lines which previously exhibited a normal karyotype. (c) A comparison of percent abnormal karyotypes broken down by the number of passages after reprogramming. This subfigure only includes results on the first karyotype performed after reprogramming. (d) A similar comparison to subfigure C examining percent abnormal cultures broken down by the number of passages since reprogramming, but exclusively on repeat karyotypes.

To establish how expansion of the iPSCs in culture may affect genetic stability, we next analyzed expanded tissue and unexpanded PBMC-iPSCs over time. About 20% of expanded tissue-derived iPSCs were observed to have abnormal karyotypes (defined as presenting greater than 1 in 20 cells with clonal aberrations) in their first G-band karyotype evaluation post-iPSC generation, which was typically between passages 1-25. Conversely, only 7.4% of unexpanded PBMC-derived iPSCs had abnormal karyotypes in their first assessment (Fig. 3b). Any iPSC line that was determined to have an abnormal karyotype in the first karyotype was not further cultured or evaluated. As such, between 2^nd^ and 4^th^ repeat karyotypes of an iPSC line were only evaluated on lines that were initially identified as cytogenetically normal in their first karyotype. Expanded iPSCs maintained a higher rate of karyotype abnormalities at 13.2%, which included lines that were normal in their first karyotypes, and significantly greater than unexpanded PBMC-iPSCs: Only 2.0 % (16 out of 814) of unexpanded PBMC-iPSCs acquired abnormal karyotypes in culture upon repeat karyotyping when compared to the proportion at first karyotypes, a significantly smaller proportion in comparison to expanded cell-derived iPSCs (Fig. 3b).

Upon analyzing the temporal relationship of the appearance of cytogenetic instability during passaging of expanded iPSCs, it appeared that the highest rate of abnormal karyotypes (first or repeat) occurred during passages 1-23 in the life of the iPSC line (Figs. 3c and d). This passage range is typically when the greatest amount of iPSC expansion occurs for any given cell line largely due to characterization and cell banking coinciding around these passage numbers. Nevertheless, this is a similar process for both expanded -iPSCs or unexpanded PBMC-iPSCs, and yet, critically, the unexpanded PBMC-derived cells reprogrammed by our new method do not display an increased disposition towards abnormal karyotypes upon extended culture or expansion.

### Unexpanded PBMC-iPSCs Acquire Less Submicroscopic Amplifications and Deletions

Since G-band karyotype is unable to detect submicroscopic genomic abnormalities (<5Mb) array comparative genomic hybridization (aCGH) microarrays were next used to determine the genome stability of a subset of fib-iPSC and PBMC-iPSC lines. While aCGH is unable to detect balanced translocations, inversions and < 20% culture mosaicism, it is can detect changes in chromosome number and copy, duplications, deletions, and unbalanced translocations and has been used in a number of iPSC lines (Spits *et al*., 2008; Elliott, Hohenstein Elliott and Kammesheidt, 2010; Martins-Taylor *et al*., 2011; Martins-Taylor and Xu, 2012). We analyzed and recorded any *de novo* copy number changes acquired in the iPSCs upon comparison with the parental fibroblast or PBMC source bio-specimen. In this analysis, amplifications and deletions acquired *de novo* in iPSCs that are not considered normal population variants (non-pathogenic and reported with population frequencies in the Database of Genomic Variants (DGV) were considered to be CNVs acquired by the reprogramming process.

Including iPSC lines with abnormal and normal G-band karyotypes, the average size of the amplification and deletions (amps/dels) detected by aCGH were significantly greater in fib-iPSC lines at 44 Mb compared to 2.1 Mb in PBMC-iPSC lines (Fig. 4a), the preponderance of which was due to fib-iPSC lines with an abnormal karyotype (events involving gain/loss of an entire chromosome) (Fig. 4b). Upon segregating the size analysis comparison between iPSC lines with normal karyotypes, the *de novo* CNVs identified were on average 2.31 Mb in fib-iPSCs and 2 Mb in PBMC-iPSC lines (Fig. 4b). Supporting this data, the average number of acquired *de novo* total CNVs in fib-iPSCs (3.7) were significantly greater (two-fold) than in PBMC-iPSCs (1.8) (Fig. 4c). Even in iPSC lines that were determined to have normal G-band karyotypes, the number of new amps/dels was greater in fib-iPSCs at 3.3 vs. PBMC-iPSCs at 1.8 (Fig. 4d). The most commonly acquired submicroscopic (0.8-1.5 Mb) *de novo* amplifications or deletions detected by aCGH were amplification of chromosome 7q31.32 or deletion of chromosomes 10q15.2-q25.1, 16p11.2, and 21p11.2-p11.1 (Fig. 4e).

**Figure 4.**
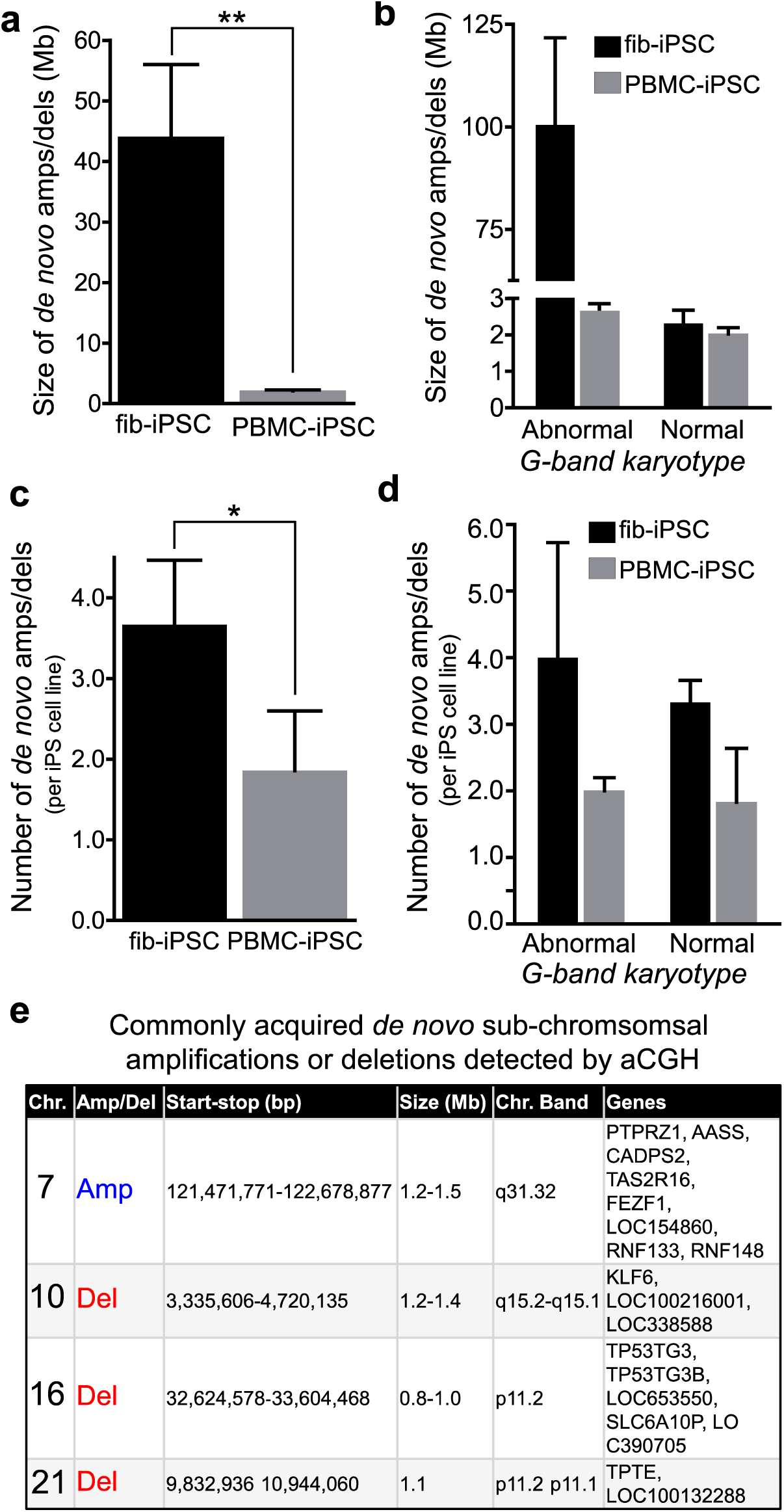
Comparative Genomic Hybridization Confirms that the PBMC-iPSCs have Relatively Small and Fewer Submicroscopic Cytogenetic Aberrations Compared to fib-iPSCs. (a) The mean size of *de novo* amplifications and deletions between fibroblast-derived iPSCs (expanded) and PBMC-derived iPSCs (unexpanded). (b) The mean size of amplifications and deletions between fibroblast and PBMC derived iPSCs when segregating between amplifications/deletions which occurred in a line with a normal G-band karyotypes vs those with an abnormal G-band karyotype. (c) The mean number of *de novo* amplifications or deletions per cell line between fibroblast and PBMC derived iPSCs. (d) Subfigure D displays the number of *de novo* amplifications and deletions when segregating out cultures which exhibited a normal G-band karyotype from those exhibiting an abnormal G-band karyotype. (e) Commonly occurring *de novo* amplifications and deletions detected by array Comparative Genomic Hybridization.

### Whole Genome Sequencing Confirms that Unexpanded PBMC-iPSCs Have a Low Mutational Burden

To detect *de novo* mutations that could be acquired during the reprogramming and culturing process three PBMC-iPSC lines were selected for whole genome sequencing. We define *de novo* variants as those that were not detected in the matched donor germline blood-derived DNA sample. DNA was extracted from whole blood collected at the time of the sample collection used to derive the PBMC-iPSC and from three clones from each PBMC-iPSC line. Each sample was sequenced to an average read depth of 84x (range 72.7x-87.5x) (Supplementary Figure 5). Single nucleotide variants (SNVs), indels and structural variants (SVs) (deletion, duplication, inversion, translocation) were identified in each sample. MuTect (Cibulskis *et al*., 2013) was used to identify *de novo* SNVs and Indels observed in only PBMC-iPSC and the consensus based workflow Parliament was used to identify SVs. We first identified variants that were observed in all three clones from a single donor, which would suggest these events are introduced during reprogramming and early in the culture/expansion of reprogrammed cells. Such variants could arise via two mechanisms; variants present in a small proportion of the germline (blood) derived sample and selected during the iPSC generation (these variants would be seen in all three of the genotyped clonal iPSC populations from each donor); variants not detected in the germline (blood) derived sample could have been introduced in the single cell(s) during clonal selection before expansion (these variants would most likely be present in all three of the reprogrammed clonal iPSCs). We identified 271 SNVs that were detected in all three clones from a single donor (79, 81, 111 for donor CS-007, CS-003 and CS-002 respectively) as candidate *de novo* variants (**Supplementary Table 5**). After further quality control to remove likely false positives (filters described in Methods) and variant visualization these were identified as false positive *de novo* variants called by the somatic variant caller (Supplementary Figure 5a). Structural variants were called using the Parliament workflow on DNAnexus, which applies a consensus approach that includes five individual software packages for calling SVs from WGS (English *et al*., 2015). SVs that were supported by >3 of the five callers were retained as high confidence SVs in each sample. We used SURVIVOR (Jeffares *et al*., 2017) to identify the SVs that were absent in the blood-derived normal and present in all three PBMC-iPSC clones from each donor (Supplementary Figure 5b). Each of the 207 SVs detected in all three clones from a single donor were visualized in IGV to confirm the genotype and identify potential variation in breakpoints for the event across the three blood-derived normal and PBMC-iPSCs. We did not confirm any *de novo* SVs present in all three clones from a single PBMC-iPSC line called by three or more of the SV calling algorithms used in the consensus SV detection pipeline. Each of the candidate *de novo* SVs was confirmed as a false positive, due to the lack of SV identification in the blood derived normal sample, where the SV was confirmed to be present, but was not called by >3 of the 5 callers. An example of such a locus is shown in Supplementary Figure 6. These results suggest that recurring structural variation is not introduced during reprogramming or culture of PBMC-iPSCs.

We then identified putative *de novo* SNVs, Indels and SVs in each clone. After applying QC filters to exclude variants within repetitive regions of the genome and retain only variants called with a base quality score >10 and an alternate allele on more than 6 reads (alternate allele fraction >0.2 and read depth >30) we identified an average of 728 putative *de novo* SNVs per clonal PBMC-iPSC line (range 481-994; **Supplementary Table 5**, **Supplementary Table 6**). Of the 6,552 SNVs and Indels identified across the 9 lines, 4.5% are annotated in dbSNP. To identify the potential function of these variants we used the Personal Cancer Genome Reporter (Nakken *et al*., 2018) to annotate each SNV with 14 annotation resources that include CIViC, VEP, ClinVar, TCGA, ICGC-PCAWG, CancerMine and OncoScore. This tool assigns variants to tiers of potential clinical and deleterious impact based on these annotations, with Tier 1 variants being those of strong clinical significance, Tier 2 variants being of potential clinical significance, Tier 3 variants being of unknown clinical significance and Tier 4 variants being coding variants with no identified clinical or deleterious impact. We did not identify Tier 1 or Tier 2 variants in any PBMC-iPSC clone (**Supplementary Table 6**). We did identify a small number of Tier 3 and Tier 4 variants per line (total of 96 SNVs across the 9 lines), and so these were each visualized in IGV for manual review. Of the 96 Tier 3 and Tier 4 SNVs identified 72 were confirmed as potential true *de novo* SNVs. A single SNV in an oncogene was identified; a missense mutation in NOTCH3, which is predicted to be deleterious by some prediction tools (ie. SIFT), but not others (ie. LRT), and to cause a change at the amino acid level (p.Met1903Arg), but is of unknown clinical significance (**Supplementary Table 7**). The mutational burden was also calculated as part of variant annotation with PCGR, and was identified as being low; 0-5 mutations/Mb, in each of the nine lines (**Supplementary Table 6**). The average mutational burden was 0.4 mutations/Mb (range 0.15-0.53) and was predominantly the result of non-coding variation. We were not able to calculate microsatellite instability or mutational signatures with the PCGR in our lines due to the very small number of putative *de novo* variants detected.

An average of 302 putative *de* novo structural variants were identified in each clone (Supplementary Figure 6a; **Supplementary Table 8**), with an average of 197 deletions, 60 duplications and 45 inversions identified per line. Most SVs were unique to a single clone and the proportions of SVs of different sizes across the clones was not different (Supplementary Figure 7; **Supplementary Table 9**). The majority of deletions and duplications were <1kb in size (85% and 75%, respectively). Inversions were more likely to be longer (>1Mb in length), but as they do not alter the dosage of DNA they are suspected to have less impact on gene expression or function than deletions or duplications.

We next analyzed these findings in the context of previously published results. We collated each mutation (SNV, Indel and SVs) previously reported in iPSCs (Table 3, **Supplementary Table 10**) (Gore, Li, Fung, Jessica E Young, *et al*., 2011; Laurent *et al*., 2011; Cheng *et al*., 2012; Bhutani *et al*., 2016; Kilpinen *et al*., 2017; Lo Sardo *et al*., 2017; Merkle *et al*., 2017). We did not identify any of our single clone 6,552 putative *de novo* SNVs and Indels that overlap with the previously reported variants. A total of 6,552 SNVs and Indels were identified across the 9 lines, of which 4.5% are annotated in dbSNP. We identified 4 genes potentially affected by low impact mutations (Tier 3 annotations by PCGR); NOTCH3, SOX17, EPHA3, ZFHX3. Notably, we identified many fewer putative *de novo* variants than previously reported methods, with 883-1249 variants (sum of SNVs, indels and SVs per line), compared to 958-7,027 reported per line using methods that rely on reprogramming of fibroblasts (D’Antonio *et al*., 2018). We also did not identify any correlation between the number of passages a line had been grown through and the number of SNVs and indels (Figure 5c) or SVs (Figure 5d; **Supplementary Table 11**). We did identify overlap of 1649 of the 2,527 *de novo* SVs we identified across the nine PBMC-iPSC lines with any one of the previously reported 12,228 SVs that could be mapped to hg19. Given that the sum of bp covered by these 12,228 SVs is more than 35 billion base pairs (35,834,452,729 bp; more than 11x the size of the human genome), this relatively high proportion of overlap (60.5%) is likely the result of the extensive number and potential large size of SVs such as whole chromosome aneuploidy previously reported.

**Figure 5.**
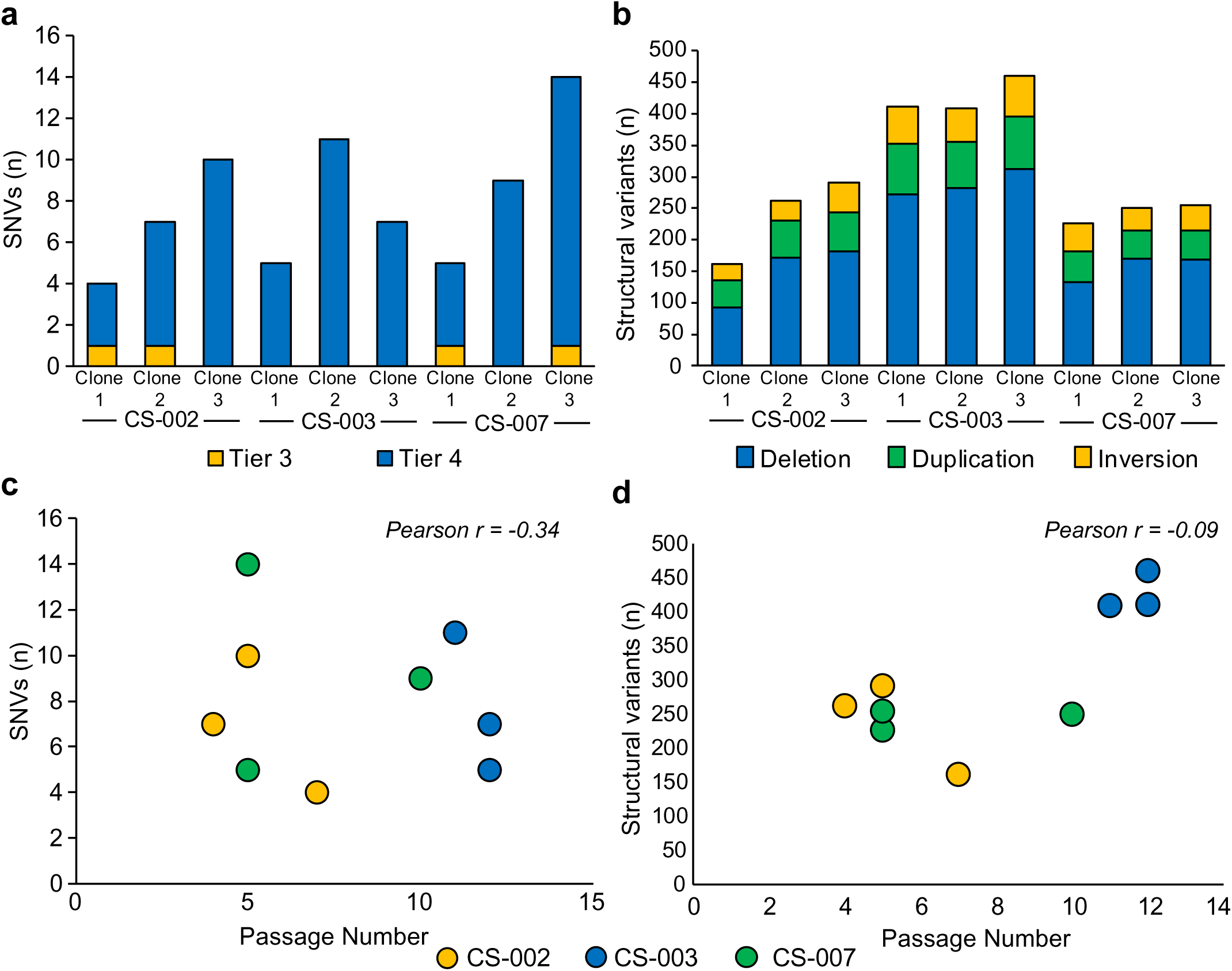
Occurrence of Single Nucleotide Variants and Structural Variants in nine cell lines. (a) The number of single nucleotide variants detected in nine profiled cell lines reprogrammed from three different parent lines. (b) The number of structural variants detected among the same profiled lines. (c) A plot of the relationship between the number of single nucleotide variants and the passage number of the line. (d) A plot of the relationship between the number of structural variants detected and the passage number of each profiled line.

**Table 3.** . **Previously reported *de novo* SNV and SV events in previously published studies of iPSC lines are not introduced during reprogramming and culture of PBMC-iPSC lines using the 5p method.** No previously reported iPSC *de novo* mutations or SVs were detected in all three clones from a single donor by whole-genome sequencing (WGS) analysis. NA=Variant identifiers or chromosome and position not available in previous publication for validation, where possible gene level intersections were performed and did not identify any shared mutations.

| Author, Year | PMID | Variant Type | N | N observed<br>PBMC-iPSC clones |
| --- | --- | --- | --- | --- |
| Laurent et al., 2011 | 21211785 | Deletion | 9707 | None |
|  |  | Duplication | 1555 | None |
| Gore et al., 2011 | 21368825 | SNV | 64 | None |
| Cheng et al., 2012 | 22385660 | SNV | 3975 | None |
| Bhutani et al., 2016 | 26892726 | SNV | 5448 | None |
|  |  | Indel | 1562 | None |
| Lo Sardo et al., 2017 | 27941802 | SNV | 863 | None |
| Merkle et al., 2017 | 28445466 | SNV | 17 | None |
| Kilpinen et al, 2017 | 28489815 | Deletion | 287 | None |
|  |  | Duplication | 783 | None |
| D'Antonio et al, 2018 | 30044985 | SNV | 39,100 | *NA |
|  |  | DNV | 2,171 | *NA |
|  |  | Indel | 2,776 | *NA |
|  |  | Deletion | 173 | None |
|  |  | Duplication | 82 | None |

## DISCUSSION

In this study, we show that human iPSCs derived by a method from unexpanded cryopreserved PBMCs (PBMC-iPSCs) are remarkably cytogenetically stable compared to iPSCs generated from other somatic cells that require cell expansion prior to reprogramming, including skin fibroblast cultures, LCLs, epithelial cells, and adipose cells. These PBMC-iPSCs reprogrammed with the 5p method also have enhanced stability rates compared to those historically reported for other pluripotent stem cell lines such as hESCs and iPSCs derived in other laboratories(Baker *et al*., 2007; Catalina *et al*., 2008; Spits *et al*., 2008; Gore, Li, Fung, Jessica E. Young, *et al*., 2011; Hussein *et al*., 2011; Laurent *et al*., 2011; Martins-Taylor *et al*., 2011; Taapken *et al*., 2011; Rebuzzini *et al*., 2015; Schlaeger *et al*., 2015). This entire study entailing episomal plasmid reprogramming of various cell types from 1,028 unique donors, identical stem cell maintenance procedures, pluripotency characterization, and karyotype analysis were performed in our laboratories and the cytogenetics core. We systematically analyzed 1,465 individual clonal PBMC-iPSC lines generated in our laboratory, with both lymphoid T cell and myeloid non-T cell populations being reliably reprogrammed, irrespective of substrate and media. Further, we demonstrated creation of stable PBMC-iPSCs on a xeno-free recombinant human recombinant L521 substrate and defined reprogramming media. Most importantly, the PBMC-iPSCs reprogrammed with the 5p method maintain karyotype stability upon long-term culture. Collectively, the combination of this improved technology and an unexpanded PBMC cell source offer both clinically compatible reprogramming methods and facile access to large numbers of patient samples.

Aneuploidy and rearrangements in chromosomes 1, 12, 13, 14 and X were the most frequent abnormalities observed in expanded source tissue-derived iPSCs generated by episomal reprogramming in our laboratory. In contrast these aberrations were rarely detected in any of the unexpanded PBMC-iPSCs we generated and analyzed. Some of these changes observed in expanded iPSCs are similar to recurrent abnormalities found in human ESCs where there are short arm of chromosome 12 and gains or losses of chromosomes 1 and X, similar to their malignant human embryonal carcinoma (EC) stem cell counterparts(Baker *et al*., 2007; Catalina *et al*., 2008; Spits *et al*., 2008; Taapken *et al*., 2011; Rebuzzini *et al*., 2015). Using aCGH microarray, the PBMC-iPSCs showed half the number of *de novo* CNVs compared to the fib-iPSCs. These *de novo* CNVs were not shared between hiPSCs and their respective parental fibroblasts or PBMCs and were likely acquired during reprogramming or iPSC expansion. The average base pair size of chromosomal change in the CNVs in fib-iPSCs was also far greater when compared to PBMC-iPSC lines, further supporting the stability of this new reprogramming method.

Increased karyotype abnormalities, coding mutations, and small genomic alterations have previously been reported in iPSCs(Gore, Li, Fung, Jessica E. Young, *et al*., 2011; Hussein *et al*., 2011; Laurent *et al*., 2011; Martins-Taylor *et al*., 2011; Taapken *et al*., 2011), likely reflecting the mutagenic consequences due to the reprogramming procedure itself. However, the bulk of these reported genomic abnormalities have been examined in iPSC lines generated from skin fibroblasts using integrating retroviral and lentiviral reprogramming methods. Aneuploidy rates and CNVs comparing different non-integrating and lentiviral reprogramming methods for low-passage iPSCs have been described(Taapken *et al*., 2011; Schlaeger *et al*., 2015). They reported an aneuploidy rate around 12-13% in low-passage iPSCs using episomal and integrating reprogramming methods, although this data predominantly includes aberration rates for iPSCs derived from expanded fibroblast cultures that were not segregated by the origin of donor cells or obtained from unexpanded cells, like PBMCs. Various factors can affect genetic stability during long-term culture and expansion of PSCs including, oxygen tension, growth supplements, growth factors, passage technique, cryopreservation and the extracellular substrate on which cells are grown(Mitalipova *et al*., 2005). There is often substantial laboratory variation in PSC culture techniques given these afore-mentioned factors; however, these factors were all tightly controlled for in our study. The Progenitor Cell Biology Consortium (www.synapse.org) identified correlations in gene expression and CNVs among key developmental and oncogenic regulators as a result of donor line stability, reprogramming technology, and cell of origin(Salomonis *et al*., 2016). However, karyotype abnormalities and line stability were not distinguished by cell of origin and their previous cell expansion history in this study, and only small numbers of abnormal fib-iPSCs were analyzed. Although methylation and transcription profiles of iPSCs segregated somewhat based on blood and fibroblast cell of origin, they could not be directly attributed to somatic cell epigenetic memory profiles. This group also observed a trend towards higher level of CNVs in lines generated using integrating reprogramming methods. Recently, another group revealed that blood cells and derived iPSCs were much closer to *bona fide* hiPSCs and ESCs in their methylation profiles (more hyper-methylated state) compared to any other parental tissue or their iPSCs that had more aberrant DNA methylation(Nishizawa *et al*., 2016), supporting our argument that a cell population that has not been previously expanded in tissue-culture, such as adult PBMCs or cord blood, could be a preferred choice for reprogramming and maintenance of cytogenetic stability at the iPSC stage.

As most SNVs are the result of replication errors(Busuttil *et al*., 2006) there is the potential for mutations to be introduced to PBMC-iPSC during passaging and expansion of the cell population. The germline and somatic mutation rates in humans have been previously estimated at 3.3 x 10^-11^ and 2.8 x 10^-7^ mutations per bp per mitosis, respectively(Milholland *et al*., 2017). These rates are considerably higher than the rate we observed in our unexpanded PBMC-iPSCs, and this could be due to several factors. Firstly, we have used PCR-free whole genome sequencing with a sequencing depth that likely reduced our false discovery rate(Fang *et al*., 2014). Secondly, we have used a stringent set of thresholds to exclude variants that could be erroneously called because of repetitive elements (particularly mis-calling SNVs within an indel) or reduced mapping quality of reads carrying the alternate allele of the acquired SNV. While our deep WGS includes only three donor PBMC-iPSC lines the lack of any acquired mutations in three clones from a single line indicate the reprogramming process is largely error-free and does not cause the introduction of mutations to the reprogrammed cells.

The mechanisms underlying the genetic stability of iPSCs derived from unexpanded PBMCs compared to iPSCs generated from expanded somatic cell types such as skin fibroblasts, LCLs, epithelial and adipose stem cells are likely to be multi-factorial. First, the bone marrow niche allows for turnover of blood more frequently (1-5 days for most cells of the PBMC lineage) than almost any other cell type of the body (BNID 101940)(Milo *et al*., 2009; Macallan *et al*., 2012). Therefore, a host blood may be considered far less “aged” than a resident dermal fibroblast, epithelial or adipose cell, which only renews over a period of months to years. Notably, the reprogramming of donor-derived skin fibroblasts, LCLs, epithelial and adipose cells, requires a significant period of *ex vivo* somatic cell expansion in tissue-culture. These factors may lead to a starting pool of cells with increased tissue culture-associated cell stress and senescence(Gorbunova *et al*., 2007; Jeyapalan *et al*., 2007; Premi *et al*., 2015), thus making susceptible to decreased DNA repair capacities(Besaratinia *et al*., 2005; Sauvaigo *et al*., 2007). It is also likely, therefore, that a culture of previously expanded somatic cells may start with some overall heterogeneity and inherent genotype instability, and low-level mosaicism could remain undetected by traditional G-band karyotype in a bulk heterogeneous cell culture prior to reprogramming. Subsequently, fib-iPSC, LCL-iPSC or epithelial-iPSC line may be isolated in which low-level karyotype abnormalities become magnified upon significant clonal proliferation and expansion. This risk is avoided in our method, as isolated PBMCs were initiated for reprogramming immediately post-isolation without any period of prior cell expansion. Lastly, the PBMC 5p reprogramming protocol contains an additional episomal plasmid where one of the reprogramming factors SV40LT minimizes PBMC cell death and promote surface attachment, which may also enhance the stability of the PBMCs during the stressful reprogramming phase for a cell. It has also been previously shown that the use SV40LT as a component of the reprogramming factor cocktail is non-transforming since the transgene does not integrate into the genome of the iPSC lines derived by episomal plasmid reprogramming(Yu *et al*., 2009), which is also supported by our data in Supplementary Figures 2 and 3. While it is possible that pre-expanded somatic cell types like fibroblasts may obtain greater genetic stability if reprogrammed with the 5p protocol, we feel that this single difference in reprogramming between the pre-expanded cells and the unexpanded PMBCs does not completely account for the vast differences in karyotype stability. Importantly, apart from the additional factor, all the pre-expanded cell types and unexpanded blood-derived iPSCs were reprogrammed using similar non-integrating episomal reprogramming methods and maintained with identical feeder-free Matrigel®/mTeSR1™ cell culture and the manual StemPro® EZPassage™ tool. Nonetheless, it would be a great advancement to the field if our novel 5p protocol facilitated reprogramming of additional cell types beyond unexpanded PBMCs and ongoing studies are assessing fibroblast reprogramming using the optimized protocol.

The first autologous human iPSC clinical trial for treating age-related macular degeneration patients sponsored by RIKEN institute was suspended because single nucleotide variations (SNVs) and CNVs were detected in a patient iPSC line, which were not detectable in the patient’s original fibroblasts. While the CNVs were all single-gene deletions, one of the SNVs was previously identified in a cancer-related somatic mutations database(Garber, 2015; Masayo Takahashi, 2016). This emphasizes that a stable karyotype is paramount for the success of clinical applications of human iPSCs. In addition, scale-up of iPSCs and derivatives for predictive toxicology and high-throughput drug screening for therapeutic discoveries requires a source of iPSCs with substantial cell stability. This optimum reprogramming method in unexpanded PBMCs will minimize effects of acquired genomic aberrations.

Together, these results highlight that optimizing techniques of reprogramming as well as the quality and the period of pre-culture prior to reprogramming are crucial for long-term iPSC line stability. This data will assist in refining the safety of iPSC-derived cell products for clinical and research discovery applications, as well as aid in greater reproducibility in iPSC disease modeling studies.

## ACKNOWLEDGMENTS

This work was supported by Cedars-Sinai Programmatic Funds (DS and CNS). We would like to thank The David and Janet Polak Foundation. We would also like to express our gratitude to Dr. Soshana Svendsen for her critical review of the manuscript. The funders had no role in study design, data collection and analysis, decision to publish, or preparation of the manuscript.

## AUTHOR CONTRIBUTIONS

Lindsay Panther: Collection and/or assembly of data, data analysis and interpretation

Loren Ornelas: Collection and/or assembly of data, data analysis and interpretation

Andrew Gross: Collection and/or assembly of data, data analysis and interpretation

Emilda Gomez: Collection and/or assembly of data

Berhan Mandefro: Collection and/or assembly of data

Anais Sahabian: Collection and/or assembly of data

Chunyan Liu: Collection and/or assembly of data

Michelle R Jones: Whole genome sequencing and genotyping array data generation and analysis of data

Benjamin Berman: Whole genome sequencing data generation and analysis of data

Clive N. Svendsen: Concept, financial support, administrative support, manuscript writing, final approval of manuscript

Dhruv Sareen: Conception and design, financial support, administrative support, collection and/or assembly of data, data analysis and interpretation, provision of study material, manuscript writing, final approval of manuscript

## DECLARATION OF INTERESTS

A US patent US 10,221,395 B2 has been granted describing this novel and efficient method for reprogramming blood to induced pluripotent stem cells. Apart from this issued patent filing the authors have declared that no other competing financial interests exist.

## ONLINE METHODS

### Ethics Statement

Human dermal fibroblasts that were obtained from the Coriell Institute for Medical Research where the Coriell Cell Repository maintains the consent and privacy of the donors. Human PBMCs were obtained from whole blood draws or some dermal fibroblasts from skin biopsies of healthy volunteers at Cedars-Sinai under the auspices of the Cedars-Sinai Medical Center Institutional Review Board (IRB) approved protocol Pro00028662 and Pro00028515. Donor-derived epithelial and adipose were isolated at Cedars-Sinai. LCLs were acquired from public repositories such as Coriell or from biobanks at Cedars-Sinai. The reprogramming and characterization of iPSC cell lines and differentiation protocols in the present study were carried out in accordance with the guidelines approved by Stem Cell Research Oversight committee (SCRO) and IRB, under the auspices of IRB-SCRO Protocols Pro00032834 (iPSC Core Repository and Stem Cell Program), Pro00024839 (Using iPS cells to develop novel tools for the treatment of SMA) and Pro00036896 (Sareen Stem Cell Program). Appropriate informed consents were obtained from all the donors. To protect donor privacy and Confidentiality, all samples were coded and de-identified in this study.

### Reprogramming of cryopreserved PBMCs

All the human peripheral blood mononuclear cells (PBMCs) and fibroblasts reported in this study were derived from multiple individuals. Approximately, 5 x 10^6^ cells per nucleofection of PBMCs were nucleofected with either plasmid mixture 4p or plasmid mixture 5p using program V-024 on the Amaxa Nucleofector 2D Device with the Amaxa Human T-cell Nucleofector® Kit. Approximately 1 x 10^6^ cells were then seeded into wells of a 6-well plate covered with mitomycin treated mouse embryonic feeder (MEF) layer or coated with 10 µg/ml Laminin-521 (L-521; BioLamina). Each episomal plasmid (Addgene) expressing 7 factors: *OCT4, SOX2*, *KLF4*, *L-MYC*, *LIN28*, *SV40LT* and *p53* shRNA (pEP4 E02S ET2K, pCXLE-hOCT3/4-shp53-F, pCXLE-hUL, pCXLE-hSK, and pCXLE-EBNA1). This method has a significant advantage over viral transduction, because exogenously introduced genes do not integrate and are instead expressed episomally in a transient fashion. Cells were plated in 2 mL of either αβ T-cell medium (X-vivo10 supplemented with 30U/ml IL-2 and 5ul/well Dynabeads Human T-activator CD3/CD28) or non-T-cell medium (αMEM supplemented with 10% FBS, 10ng/ml IL-3, 10ng/ml IL-6, 10ng/ml G-CSF and 10ng/ml GM-CSF). Two days after nucleofection an equal amount of Primate ESC medium (ReproCell) containing 5 ng/ml bFGF (for MEF condition) or E7 medium (for L-521 condition) was added to the wells without aspirating the previous medium. Beginning on day four, the medium was gently aspirated from each well and 2ml of the appropriate fresh reprogramming media was added to each well. Medium was replaced every other day. At approximately day 18 post nucleofection, individual colonies were observed in all wells of each condition. At approximately day 25 post nucleofection, individual colonies were isolated and sub-cloned into 1 well of 12-well plate containing the appropriate substrate and medium. These nucleofected cells were plated on feeder-independent BD Matrigel™ growth factor-reduced Matrix (Corning/BD Biosciences, #354230). All cultures were maintained at 20% O_2_ during the reprogramming process. Individual PBMC-iPSC colonies with ES/iPSC-like morphology appeared between day 25-32 and those with best morphology were mechanically isolated, transferred onto 12-well plates with fresh Matrigel™ Matrix, and maintained in mTeSR®1 medium. The iPSC clones were further expanded and scaled up for further analysis. All the fibroblasts were reprogrammed with the 3p reprogramming plasmid mixture as described previously(Okita *et al*., 2011; Sareen *et al*., 2013).

### Plasmid Mixture (3p)(Okita *et al*., 2011)

| Plasmid | Reprogramming Factors | Amount |
| --- | --- | --- |
| pCXLE-hOCT3/4-shp53 | *OCT3/4 (POU5F1)*, shRNA to *TP53* | 0.83 µg |
| pCXLE-hSK | *SOX2*, *KLF4* | 0.83 µg |
| pCXLE-UL | *LIN28*, *L-MYC* | 0.83 µg |

### Plasmid Mixture (4p)(Okita *et al*., 2013)

| Plasmid | Reprogramming Factors | Amount |
| --- | --- | --- |
| pCXLE-hOCT3/4-shp53 | *OCT3/4 (POU5F1)*, shRNA to *TP53* | 0.83 µg |
| pCXLE-hSK | *SOX2*, *KLF4* | 0.83 µg |
| pCXLE-UL | *LIN28*, *L-MYC* | 0.83 µg |
| pCXLE-EBNA1 | *EBNA1* | 0.5 µg |

### Plasmid Mixture (5p)

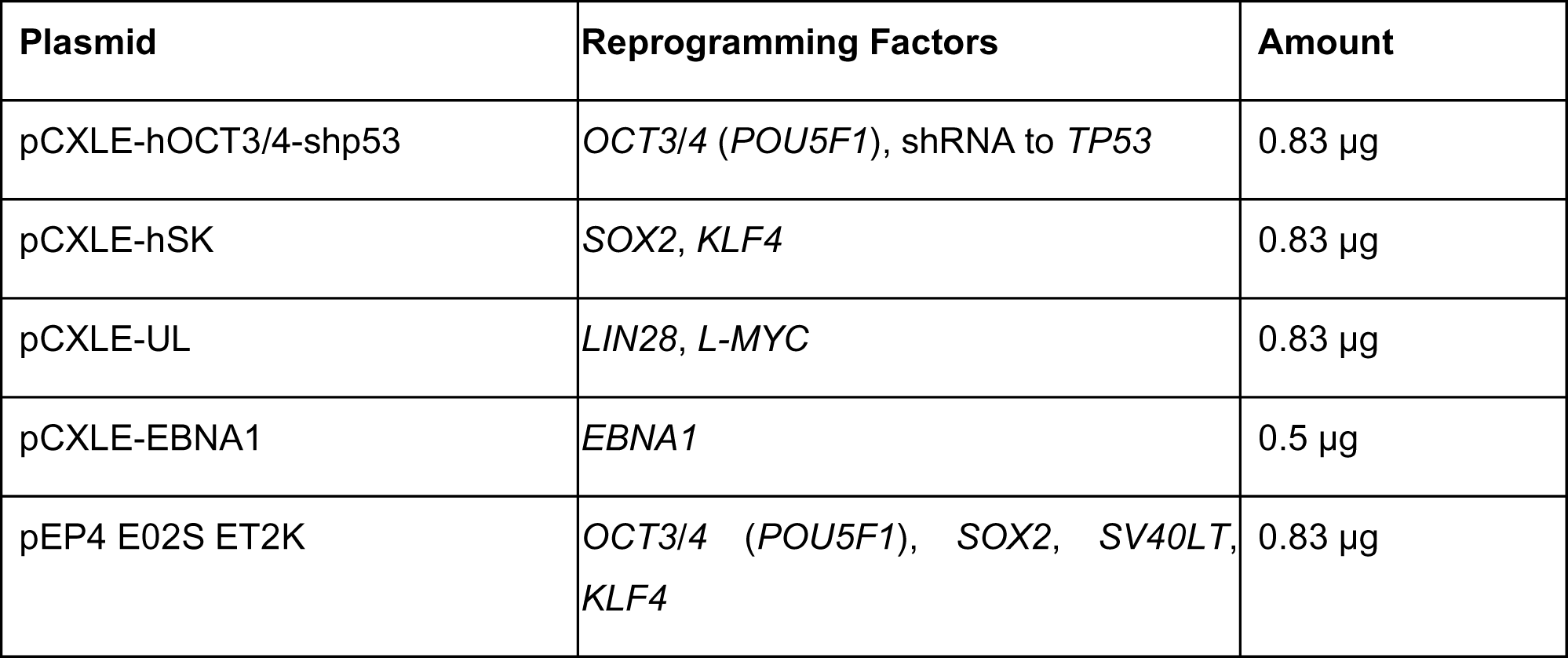

### Cell Culture

PBMCs were purified by BD Vacutainer CPT (REF 362761; BD Biosciences-US) per the manufacturer’s instructions. Isolated PBMCs were frozen in a 1:1 mixture of human plasma and Cryostor CS 10. Human induced pluripotent stem cells (iPSCs) that were generated on MEFs were slowly adapted to feeder-free culture conditions (growth factor reduced Matrigel and mTeSR1) at approximately p3-p4. Human iPSCs that were generated on L-521 were maintained on 10 µg/ml L-521 (BioLamina) in home-made E8 culture medium. iPSCs were passaged every 5-7 days using the StemPro EZ Passaging Tool (Life Technologies) or Versene solution (Life Technologies). Every cell line in the iPSC core cultured is tested monthly for mycoplasma using the MycoAlert™ PLUS Mycoplasma Detection Kit (Lonza; LT07-710). Our annual rate of mycoplasma contamination is less than 0.2% and only mycoplasma-free cell lines were used in this study for analysis.

### Characterization of iPSC Clones

The iPSC clones were subjected to various characterization assays including alkaline phosphatase (AP) staining (Stemgent), immunocytochemistry staining for the presence of pluripotency markers (SSEA4, OCT4, Tra -1-81, Tra-1-60, Nanog, and Sox2; see **Supplementary Table 12** for manufacturer and catalog information), Illumina gene-chip expression and bioinformatics assay (PluriTest), RT-qPCR to confirm endogenous expression of pluripotency and confirm the lack of exogenous gene expression, G-band karyotype, and EB formation. T-cell lineage was confirmed via PCR-based detection of the TRB rearrangement using the TCRB Gene Clonality Assay (InVivoScribe) per the manufacturer’s instructions. The iPSC lines once reprogrammed are authenticated using cell line identity assay by short tandem repeat (STR) profiling and compared to parental tissue source (PBMCs or fibroblasts). For STR profiling, the iPSC lines were authenticated by contracting with IDEXX Laboratories, Inc. and use of their CellCheck 9 service. This test allows provides a human 9 STR marker profile and for interspecies contamination check for human, mouse, rat, African green monkey and Chinese hamster cells. The identity of all cell lines validated is matched to tissue source annually and at the time of generation of the distribution cell bank. Certificate of Analysis of all iPSC lines used in this manuscript are available upon request.

### Alkaline phosphatase staining

Alkaline Phosphatase staining was performed using the Alkaline Phosphatase Staining Kit II (Stemgent, Cat no. 00-0055) according to the manufacturer’s instructions.

### Immunohisto/cytochemistry

PBMC-iPSCs, fib-iPSCs or differentiated cells were plated on glass coverslips or optical-bottom 96-well plates (Thermo, # 165305) and subsequently fixed in 4% paraformaldehyde. All cells were blocked in 5-10% goat or donkey serum (Millipore) with 0.1% Triton X-100 (Bio-Rad) and incubated with primary antibodies (**Supplementary Table 12**) either for either 3 hours at room temperature or overnight at 4°C and subsequently washed with PBS. Cells were then rinsed in PBS with 0.1% Tween-20 and incubated in species-specific AF488 or AF594-conjugated secondary antibodies followed by Hoechst 33258 (0.5 µg/mL; Sigma) to counterstain nuclei. Cells were imaged using Nikon/Leica microscopes or on Image Express Micro high-content imaging system (Molecular Devices).

### G-Band Karyotype

Human iPSCs were incubated in Colcemid (100 ng/mL; Life Technologies) for 30 minutes at 37°C and then dissociated using TrypLE for 10 minutes. They were then washed in phosphate buffered saline (PBS) and incubated at 37°C in 5mL hypotonic solution (1g KCl, 1g Na Citrate in 400mL water) for 30 minutes. The cells were centrifuged for 2.5 minutes at 1500 RPM and resuspended in fixative (methanol: acetic acid, 3:1) at room temperature for 5 minutes. This was repeated twice, and finally cells were resuspended in 500 µl of fixative solution and submitted to the Cedars-Sinai Clinical Cytogenetics Core for G-Band karyotyping. Analysis includes examination of chromosomes in a minimum of 20 cells per culture. Only abnormalities that met the ISCN 2013 definition of clonality were included in this manuscript dataset(Simons, Shaffer and Hastings, 2013). All normal and abnormal karyotypes were transcribed into a table format. The chromosomal ideograms were then plotted in R using the karyoploteR Bioconductor package [[REF: Bernat & Sera 2017]].To accurately reflect the frequency of chromosomal rearrangement, the bar widths for each ideogram were scaled in proportion to the number of samples examined. As such, the bars displaying abnormalities found in PBMC-iPSC lines are 3.6 times the width of bars depicting abnormalities found in fib-iPSC lines to account for the fibroblast ideogram portraying the number of abnormalities found in a sample set that was 3.6 times larger than the PBMC set. Pie charts of the fraction of normal and abnormal lines were also plotted in R using ggplot2 [[REF: Wickman]] and <u>scatterpie</u>. All analysis scripts can be found at <u>github.com/andrewrgross/</u> under the /E427_karyotype_data_processing repository.

### Copy Number Variations with Array Comparative Genomic Hybridization (aCGH)

Array CGH is a high-resolution karyotype analysis solution for the detection of unbalanced structural and numerical chromosomal alterations with high-throughput capabilities. High quality genomic DNA was isolated from primary dermal fibroblasts or isolated PBMCs and their reprogrammed iPSCs. Quality of genomic DNA as determined by UV spec. (NanoVue), fluorometer (Qubit) and Agarose Gel analysis. The samples were analyzed for CNVs using the Agilent 60K Standard aCGH platform suitable for human stem cell and cancer cell lines by Cell Line Genetics (Madison, WI). Comparison of iPSC lines to their parental donor cell of origin yielded *de novo* copy-number variation (gains/amplifications or losses/deletions) for the indicated loci.

### PluriTest(Müller *et al*., 2011)

Total RNA was isolated using the RNeasy Mini Kit (Qiagen) and subsequently run on a Human HT-12 v4 Expression BeadChip Kit (Illumina). The raw data file (idat file) was subsequently uploaded onto the Pluritest widget online (www.pluritest.org).

### Quantitative RT-PCR (RTq-PCR)

Total RNA was isolated using the RNeasy Mini Kit (Qiagen), and 1 ug of RNA was used to make cDNA using the transcription system (Promega). RTq-PCR was performed using specific primer sequences (**Supplementary Table 13**) under standard conditions. “CDS” indicates that primers designed for the coding sequence measured expression of the total endogenous gene expression only, whereas “Pla” indicates that primers designed for the plasmid transgene expression only. Data are represented as mean ± SEM

### TCRG chain rearrangement assay

Genomic DNA (350ng) was harvested from all cell lines using the MasterPure DNA Purification Kit (Epicenter Biotechnologies). An embryonic stem cell line (H9) was used as a negative control. Primer sets that recognize the three framework regions in the heavy chain locus of the IgH gene were obtained from InVivoScribe Technologies (Cat no.11010010, San Diego, CA) and the PCR was carried out as per the manufacturer’s protocol.

### Episomal plasmid related gene analysis

Genomic DNA (400ng) was harvested from all cell lines and an embryonic stem cell line (H9) was used a negative control. Primers that recognize EBNA-1, along with GAPDH, which was used as a housekeeping gene, were included in this study. PCR was run for 35 cycles at 95°C for 30 sec, 60°C for 30 s, and 72°C for 30 s.

### Neuronal differentiation from iPSCs

The human fibroblast- and PBMC-iPSCs were seeded in 6-well Matrigel-coated plates. The iPSCs were grown to near confluence under normal maintenance conditions before the start of the differentiation. The next day neuronal differentiation was initiated by neuroectoderm differentiation by dual SMAD and GSK-3beta inhibition using LDN193189 (0.2 μM, Cayman), SB431542 (10 μM, Cayman) and CHIR99021 (3 μM, Cayman) this treatment is carried on for 6 days in Iscove’s modified Dulbecco’s medium (IMDM) / F12 (1:1) media containing non-essential amino acids (NEAA; 1%), B27 (2%), and N2 (1%). After 6 days differentiating cells were then gently lifted by accutase treatment for 5 min at 37°C. Cells at a density of 7.5 × 10^5^ were subsequently placed in a 6-well plate or 1 × 10^4^ cells were seeded in a 96-well plate in above neural differentiation medium with the addition of 0.1 μM all-trans retinoic acid and 1 μM sonic hedgehog agonist (SAG). Fresh media was changed every other day. At day 12 of differentiation media was changed to terminal differentiation (IMDM) / F12 (1:1) media supplemented with NEAA (1%), B27 (2%), N2 (1%), compound E (0.1 μM; Calbiochem), DAPT (2.5 μM; Cayman), retinoic acid (0.5 μM) all-trans, SAG (0.1 μM), ascorbic acid (200 ng/ml), dibutyryl cyclic adenosine monophosphate (1μM), brain-derived neurotrophic factor (10 ng/ml), and glial cell line–derived neurotrophic factor (10 ng/ml). Differentiating neurons were fixed at day 18 of differentiation and analyzed.

### Antibody reagents

Specificity of all commercial monoclonal and polyclonal antibodies for many of the stem cells and neuronal markers have already been validated in our laboratory (**Supplementary Table 12**). We have used well-validated antibodies that have been extensively published previously in the literature.

### Whole Genome Sequencing

Genomic DNA was extracted from whole blood or cultured PBMC-iPSCs for whole genome analysis. 1.5 µg of genomic DNA used to generate two sequencing libraries (750ng per library) at Fulgent Therapeutics. Sample quality was confirmed using the Qubit (which detects only dsDNA) and by running the samples on a 1.5% agarose gel for 1hr. Sequencing libraries were prepared using the Illumina TruSeq DNA PCR free Library Preparation Kit (Illumina, San Diego, CA) according to manufacturer’s instructions and each library was sequenced on the Illumina HiSeqX in 2×150bp format. FastQC (https://www.bioinformatics.babraham.ac.uk/projects/fastqc/), MultiQC(Ewels *et al*., 2016) and Picard Tools (http://broadinstitute.github.io/picard/) were used to perform QC of the data and generate metrics for data analysis. Alignment and variant calling were performed on the DNANexus platform (Mountain View, CA) using BWA -mem to align to the reference genome build hg38 and GATKv3 including Indel realignment and base score recalibration. To identify *de novo* variants observed in only the PBMC-iPSC paired normal/germline (blood) and somatic (PBMC-iPSC) samples were analyzed using MuTect v1.1.7(Cibulskis *et al*., 2013).

SNVs and indels identified as somatic (genotype in PBMC-iPSC differing from the blood derived sample) in all three clones from a single donor sample were visualized in the Integrated Genomics Viewer (IGV) (Thorvaldsdóttir, Robinson and Mesirov, 2013). Variants were excluded if minor allele frequency <0.06 was observed in combination with low mapping quality, reads overlapping the variant with paired end reads with mates on a difference chromosome that was not identified as a translocation by the structural variant calling pipeline, a region rich in repeats, coverage <50% of average coverage across the genome, variants identified within an indel where complex alleles of the indel could explain the presence of the alternate allele at the identified position, or additional variants identified with alternate alleles present only on the same reads. SNVs and indels identified in each of the nine PBMC-iPSC lines using rthe default MuTect variant calling parameters (n=16,194) were further filtered to reduce the number of false positive variant calls. Using bcftools v1.9 (Li, 2011) variants were retained if they were flagged as passing MuTect internal quality control metrics, plus if their alternate allele was present on more than 6 reads (combination of read depth >30 and alternate allele fraction greater than 0.2) and had a base quality score >10. Finally, variants identified in highly repetitive regions were removed using the RepeatMasker (Smit, AFA, Hubley, R & Green, P. *RepeatMasker Open-4.0*, 2015 www.repeatmasker.org database. Putative *de novo* variants identified in our nine PBMC-iPSC lines were intersected with previously reported SNVs and indels from the literature using UCSC Genome Browser Table Browser intersection tool after mapping chromosome and position of each variant to hg19.

Structural variants were identified in all samples using the Parliament 0.1.4 pipeline on the DNA Nexus platform and structural variants unique to PBMC-iPSC identified. This pipeline uses five callers; LUMPY (Layer *et al*., 2014), Manta (Chen *et al*., 2016), DELLY (Rausch *et al*., 2012), BreakDancer (Chen *et al*., 2009), and CNVnator (Abyzov *et al*., 2011) to generate a set of candidate structural variants. The Parliament package (English *et al*., 2015) and SURVIVOR tool (Jeffares *et al*., 2017) then identify structural variants called by multiple individual tools, and the SVTYPER package (www.github.com/hall-lab/sv-pipeline) utilizes breakpoint evidence from both read depth and genotype to produce a likely structural event using a maximum likelihood Bayesian classification algorithm. SV events with supporting breakpoint evidence from 4+ of the five SV calling programs were retained for analysis and comparison between the blood-derived and PBMC-iPSC derived samples (**Supplementary Table 14**). Variant annotation (inference of functional consequences of variants) was performed using the Personal Cancer Genome Reporter (Nakken *et al*., 2018) and variant annotation databases used included GENCODE (Frankish *et al*., 2019), dbNSFP (Liu *et al*., 2016), Pfam (El-Gebali *et al*., 2019), TCGA (Weinstein *et al*., 2013), ICGC-PCAWG (Goldman *et al*., 2020), TCGA-PCDM (Bailey *et al*., 2018), UniProtKB (Breuza *et al*., 2016), CORUM (Ruepp *et al*., 2010), gnomAD (Karczewski *et al*., 2020), dbSNP (Sherry *et al*., 2001), 1000Genomes (Auton *et al*., 2015), DisGenet (Piñero *et al*., 2017), DoCM (Ainscough *et al*., 2016), CancerHotspots (Chang *et al*., 2018), ClinVar (Landrum *et al*., 2014), CancerMine (Lever *et al*., 2019), DiseaseOntology (Kibbe *et al*., 2015), OncoScore (Piazza *et al*., 2017), OpenTargetPlatform (Carvalho-Silva *et al*., 2019), DGIdb (Cotto *et al*., 2018), ChEMBL (Gaulton *et al*., 2017), CIViC(Griffith *et al*., 2017), CBMDB (Tamborero *et al*., 2018), ClinicalTrials.gov, ClinVar (Landrum *et al*., 2014), COSMIC (Forbes *et al*., 2017), and PolyPhen-2 (Adzhubei *et al*., 2010).

### Statistical Analysis

In this study, we report cytogenetic aberrations based upon G-band karyotype analysis for every iPSC line ever used or generated in the iPSC Core since 2011. G-band karyotype analysis was performed a total of 2,403 times (fibroblasts: 358; LCL: 122; epithelial: 42; adipose 4; PBMC:1877) on adipose 4; PBMC:1,877) on 1,465 human iPSC cultures (fibroblasts: 265; LCL: 91; epithelial: 42; adipose 4; PBMC:1,065) derived from 1,028 unique donors (fibroblasts: 98; LCL: 54; epithelial: 15; adipose 1; PBMC: 860) multiple laboratories or public repositories. This provided a total of 526 karyotypes from lines expanded prior to reprogramming and 1,877 lines from lines that were unexpanded prior to reprogramming.

All data are represented as mean ± S.D. *p* values < 0.05 were considered significant – * *p* ≤ 0.05, ** *p* ≤ 0.01, *** *p* ≤ 0.001 and **** *p* ≤ 0.0001. All statistical analyses were performed using R, GraphPad Prism, <u>In-Silico Statistical Calculator</u> and <u>Social Science Statistics.</u> All analyses were conducted using student’s unpaired t-test (with or without Welch’s correction) in Figures 1c, 4a and 4c or two-proportion z-test (also called two population proportions) in Figures 1b and 3.

The *z*-test is the assessment of proportions is used to investigate whether two populations or groups differ significantly in proportion – for example, whether there is a difference in the proportions of fib-iPSC or PBMC-iPSC groups. The significance level was set at 0.05 and we used a two-tailed (two-sided) hypothesis. The two requirements for a two-proportion *z*-test is that (1) a random sample of each of the population groups to be compared and (2) categorical data is used, for example, abnormal karyotype or normal karyotype.

The test statistic for testing the difference in *z*-test two population proportions, that is, for testing the null hypothesis

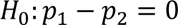

is:

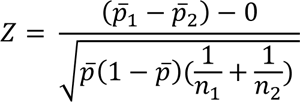

where:

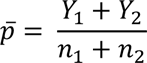

the proportion of “successes” in the two samples combined.

*p*_1_ is the proportion from the first population (fib-iPSC group)

*p*_2_ the proportion from the second population (PBMC-iPSC group)

The null hypothesis tends to be that there is no difference (zero) between the two population proportions.

## SUPPLEMENTARY FIGURES

**Supplementary Figure 1:**
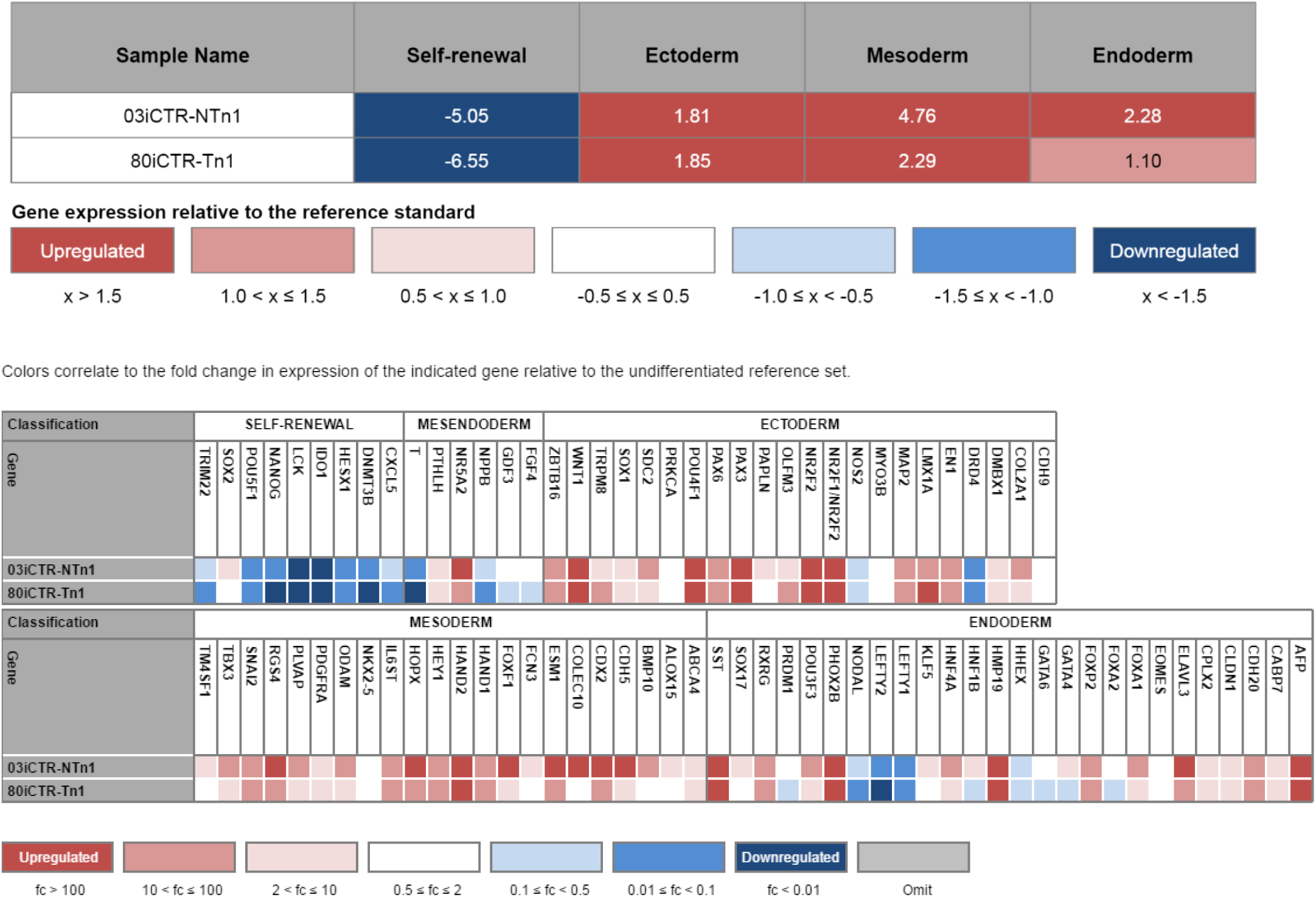
Tri-lineage potential of PBMC-derived iPSCs. TaqMan hPSC Scorecard table showing the tri-lineage potential of the representative PBMC-derived iPSCs generated in our laboratory. Expression of selected genes in four groups (self-renewal/pluripotency, ectoderm, endoderm, and mesoderm) is compared for spontaneous *in vitro* differentiation of embryoid bodies (EBs) derived from each PBMC-iPSC line between 14-18 days post-iPSC stage. Negative scores for self-renewal reflect the differentiating status of EBs, while positive scores for each of the 3 germ layers shows relatively equal propensity of a PBMC-derived iPSC line to generate the ectoderm, mesoderm and endoderm without directed differentiation.

**Supplementary Figure 2:**
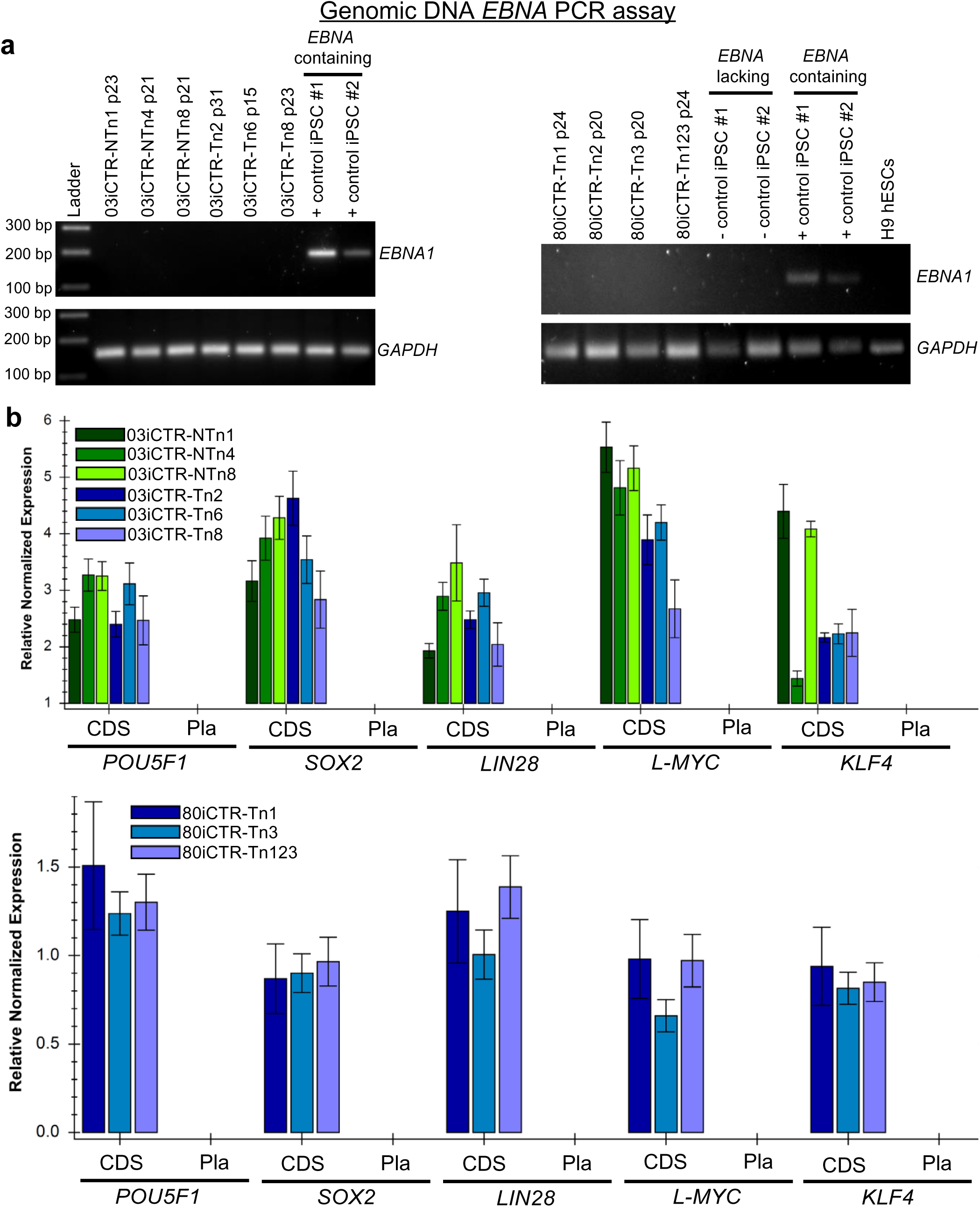
PBMC-iPSCs reprogrammed with episomal plasmids are transgene-free. a, Lack of plasmid-based *EBNA1* gene presence in the genomic DNA of the PBMC-iPSCs. **b,** Relative normalized gene expression measured by quantitative RT-PCR analyses using primers detecting endogenous *POU5F1 (OCT4)*, *SOX2*, *LIN28*, *L-MYC*, and *KLF4* expression (coding DNA sequence, CDS), as well as plasmid-derived expression (Pla), which is undetectable in the PBMC-iPSC lines.

**Supplementary Figure 3:**
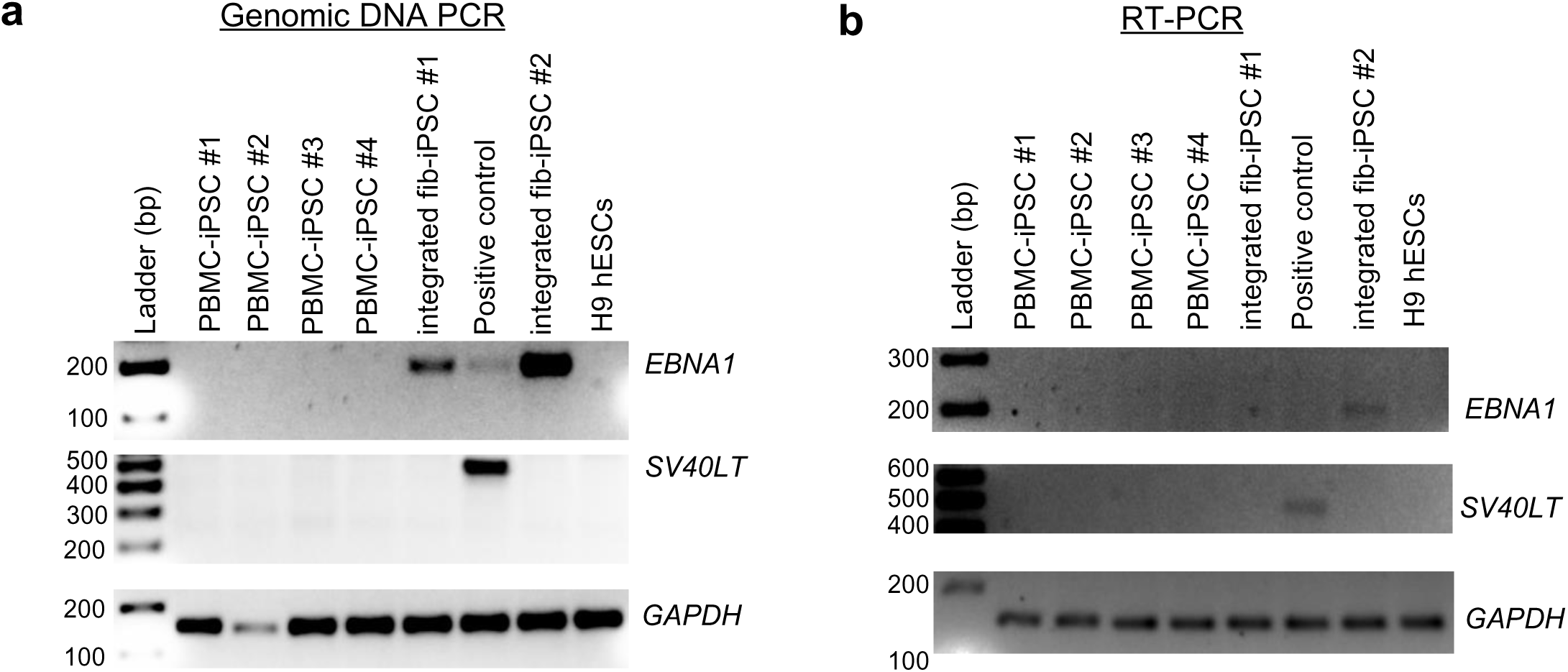
PBMC-iPSCs Reprogrammed with 5p Method are Free of *SV40LT* Gene Integration and Expression. a, Lack of plasmid-based *EBNA1* and *SV40LT* gene presence in the genomic and episomal DNA isolated from PBMC-iPSCs, H9-hESCs, integrated iPSC control lines and a positive control (nucleofected with 5p in PBMCS for 30 days). **b,** Gene expression measured by quantitative RT-PCR analyses in the same cell lines using primers detecting expression for *EBNA1, SV40LT* and *GAPDH. SV40LT* gene expression is undetectable in the reprogrammed PBMC-iPSC lines, when compared to the nucleofected positive control. 32 PCR cycles were used for all primer sets.

**Supplementary Figure 4:**
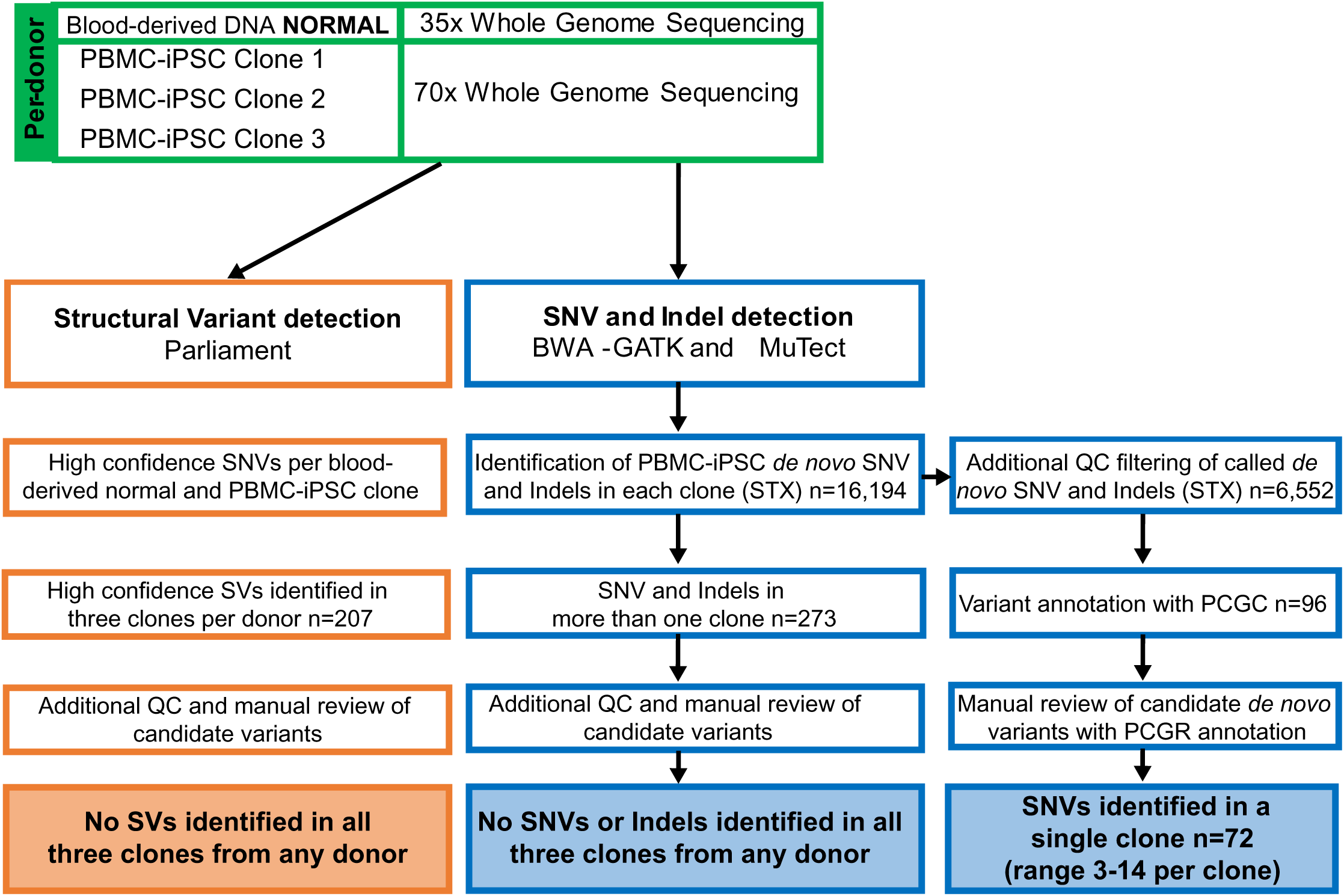
Whole genome sequencing data generation and analysis workflow.

**Supplementary Figure 5:**
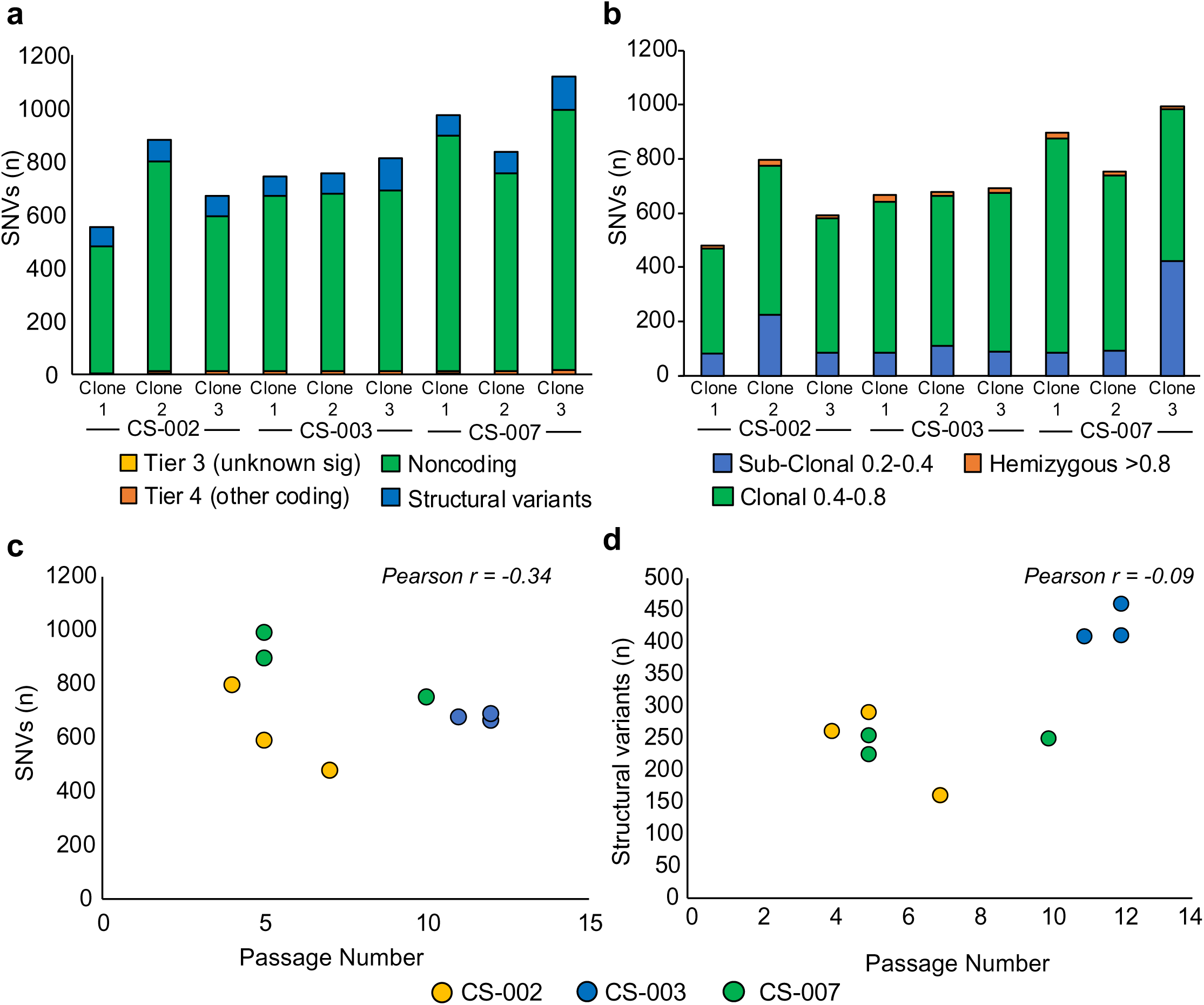
iPSC reprogramming of PBMCs with the 5p method does not introduce deleterious mutations. a, The number of de novo SNVs that are predicted to be functional (of unknownsignificance or other coding variants) or non-coding, and structural variant per clone. b, Clonality of identifiedde novo SNVs per clone. c, No correlation was identified between passage number and the number of SNVmutations identified in each clone. d, No correlation was identified between passage number and the number of SVs identified in each clone. Panels a-c include all the SNVs that made it through filtering (includes non-coding SNVs that were not visually inspected for QC).

**Supplementary Figure 6:**
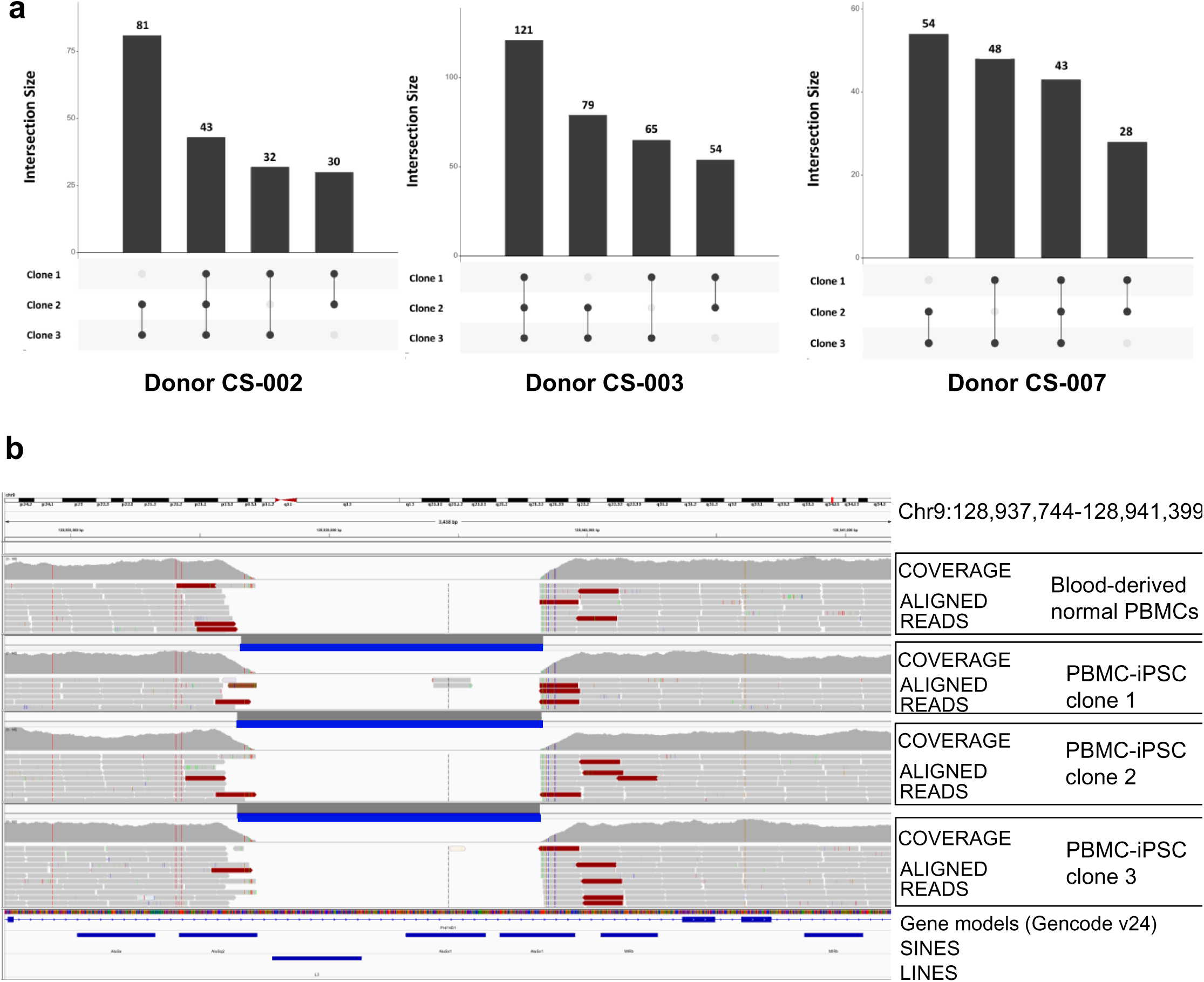
Results from analysis of Structural Variants in PBMC-iPSC Clones. a, UpsetR plots showing the number of structural variants identified in PBMC-iPSC clones from each donor. A small number of SVs unique to the PBMC-iPSC clones (not called in the blood-derived normal DNA from each donor) were identified in more than one clone. All SVs identified in all three PBMC-iPSC clones were then flagged for visualization with pre-filtered genotype call files (.vcf) and the aligned reads (.bam) used to make structural variant calls. All SVs identified as unique to PBMC-iPSC clones and observed in all three clones from each donor were confirmed to be false positives, where the event was in fact observed in the blood derived normal PBMCs, but didn’t meet our threshold for calling high confidence events (supported by breakpoint evidence from >4 callers with SVTyper). **b,** example of visualization of a false positive PBMC-iPSC structural variant. Track 1 shows the coverage and aligned reads for blood-derived normal DNA from donor CS-007, but no high confidence call for this deletion was reported (supporting evidence for this event came from 3 of the 5 callers). Track 2-4 shows the identified deletion in each of the three PBMC-iPSC clones from this donor. Track 5 shows reference genome sequence, gene models from Gencode v24 and two repeat element tracks (SINES and LINES) used to identify likely false positives.

**Supplementary Figure 7:**
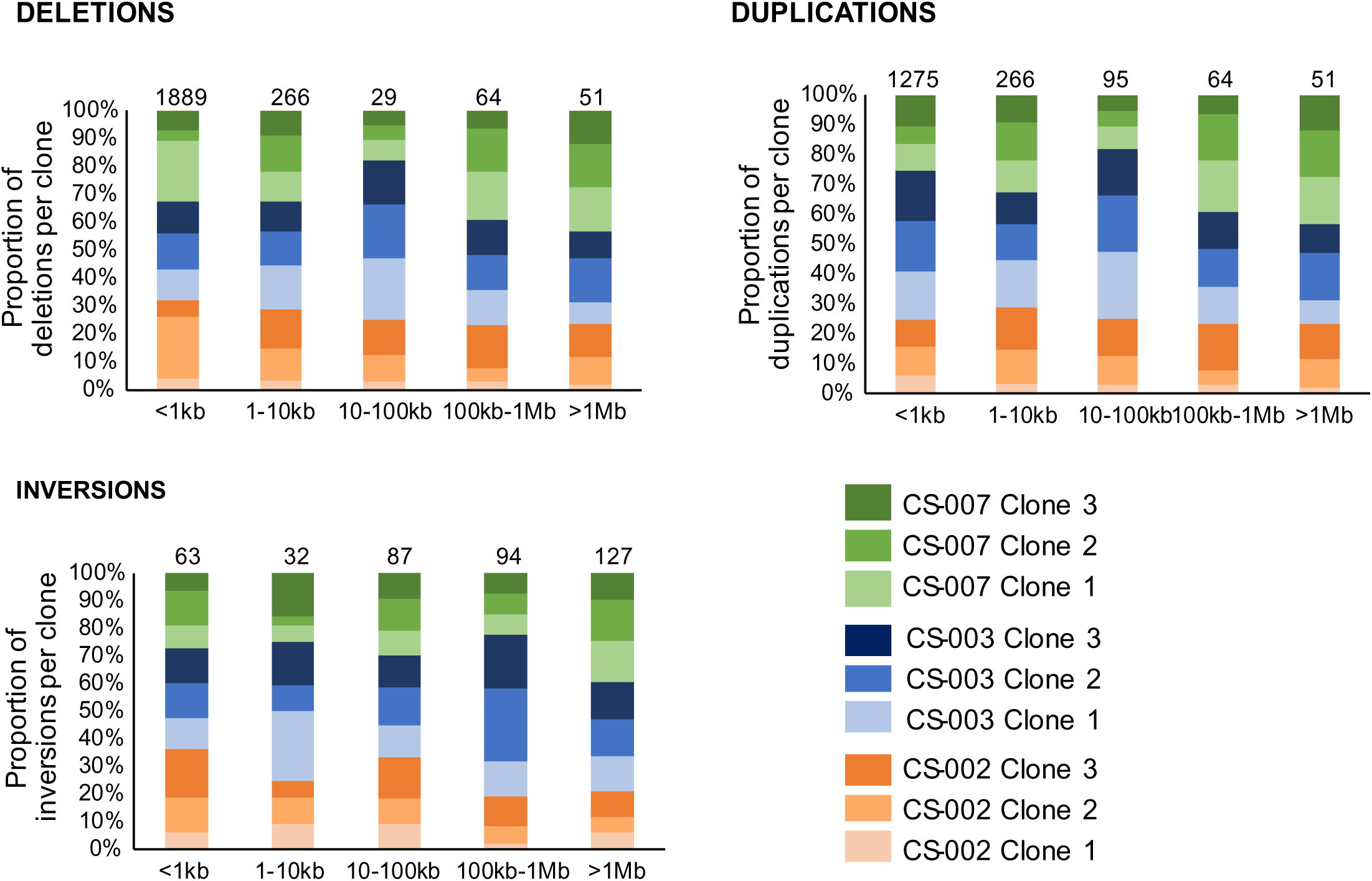
Number of structural variants by variant size identified in each PBMC-iPSC clone.

## Notes

https://docs.google.com/spreadsheets/d/1rboYasZNdsZ_tgMyhGQlqcxsH8CVBDhK/edit?usp=sharing&ouid=110828706727380725279&rtpof=true&sd=true

